# No evidence of theory of mind reasoning in the human language network

**DOI:** 10.1101/2022.07.18.500516

**Authors:** Cory Shain, Alexander Paunov, Xuanyi Chen, Benjamin Lipkin, Evelina Fedorenko

## Abstract

Language comprehension and the ability to infer others’ thoughts (theory of mind, ToM) are interrelated during development and language use. However, neural evidence that bears on the relationship between language and ToM mechanisms is mixed. Although robust dissociations have been reported in brain disorders, brain activations for contrasts that target language and ToM bear similarities, and some have reported overlap (Deen et al., 2015). We take another look at the language-ToM relationship by evaluating the response of the language network (Fedorenko et al., 2010), as measured with fMRI, to verbal and non-verbal ToM across 151 participants. Individual-subject analyses reveal that all core language regions respond more strongly when participants read vignettes about false beliefs compared to the control vignettes. However, we show that these differences are largely due to linguistic confounds, and no such effects appear in a non-verbal ToM task. These results argue against cognitive and neural overlap between language processing and ToM. In exploratory analyses, we find responses to social processing in the “periphery” of the language network—right hemisphere homotopes of core language areas and areas in bilateral angular gyri—but these responses are not selectively ToM-related and may reflect general visual semantic processing.

## Introduction

Everyday social interactions regularly involve an intricate coordination between language use on the one hand and reasoning about others’ mental states (theory of mind, or ToM) on the other. For example, to understand the communicative intent behind an utterance (e.g., *Nice outfit*), we often rely on inferences about the speaker’s state of mind (e.g., whether they are likely to have a positive assessment of your outfit, and thus whether the utterance was likely sincere or sarcastic). In addition to this kind of ToM-based pragmatic reasoning needed to infer implicit meanings from utterances (e.g., Grice, 1975; Sperber & Wilson, 1987; Winner et al., 1998; Champagne-Lavau & Joanette, 2009; Roberts, 2012), linguistic representations may be critical to the development of ToM (e.g., Astington & Jenkins, 1999; Peterson & Siegal, 2000; Hale & Tager-Flusberg, 2003; Ruffman et al., 2003; Astington & Baird, 2005; Slade & Ruffman, 2005; Miller, 2006; de Villiers & de Villiers, 2014; Richardson et al., 2020). There is thus reason to suspect a close cognitive and neural connection between language and ToM processing, and perhaps even overlap between the neural resources that support both kinds of skills.

However, neuroscientific evidence that bears on the relationship between language and ToM paints a complex picture. On the one hand, at least some evidence indicates that language and ToM rely on distinct cognitive and neural mechanisms. In particular, ToM reasoning abilities can be preserved in cases of linguistic deficits (e.g., in aphasia; e.g., Dronkers et al., 1998; Varley et al., 2001; Apperly et al., 2006; Willems et al., 2011), and at least some aspects of language can be preserved when social abilities are impaired (e.g., in some individuals with autism spectrum disorders; e.g., Tager-Flusberg et al., 2005; Diehl et al., 2006). Furthermore, the core brain areas that have been linked to language vs. ToM appear to be distinct. Language processing recruits a left-lateralized network of lateral frontal and temporal areas (e.g., Binder et al., 1997; Fedorenko et al., 2010), whereas social cognitive processing, including ToM/mentalizing, recruits bilateral (though more strongly present in the right hemisphere) areas at the junction of temporal and parietal cortex along with frontal and parietal cortical midline regions (e.g., Fletcher et al., 1995; Castelli et al., 2000; Gallagher et al., 2000; Vogeley et al., 2001; Ruby & Decety, 2003; Saxe & Kanwisher, 2003, *inter alia*). These sets of areas also dissociate during naturalistic cognition: they show strong within-network correlations and weaker correlations among pairs of brain regions that straddle network boundaries (e.g., Paunov et al., 2019; Braga et al., 2020) and ‘track’ different aspects of naturalistic stimuli (Paunov et al., 2022). On the other hand, whole-brain activation landscapes for contrasts that target language processing and those that target ToM bear similarities (**Figure 1**). Further, Deen et al. (2015; cf. Koster-Hale & Saxe, 2013) examined responses to language and ToM using individual-subject analyses and reported partial overlap between language and ToM areas in the posterior temporal lobe and angular gyrus. But Deen et al.’s study used a ToM contrast based on verbal vignettes that could have linguistic differences, making these findings difficult to interpret.

**Figure 1:**
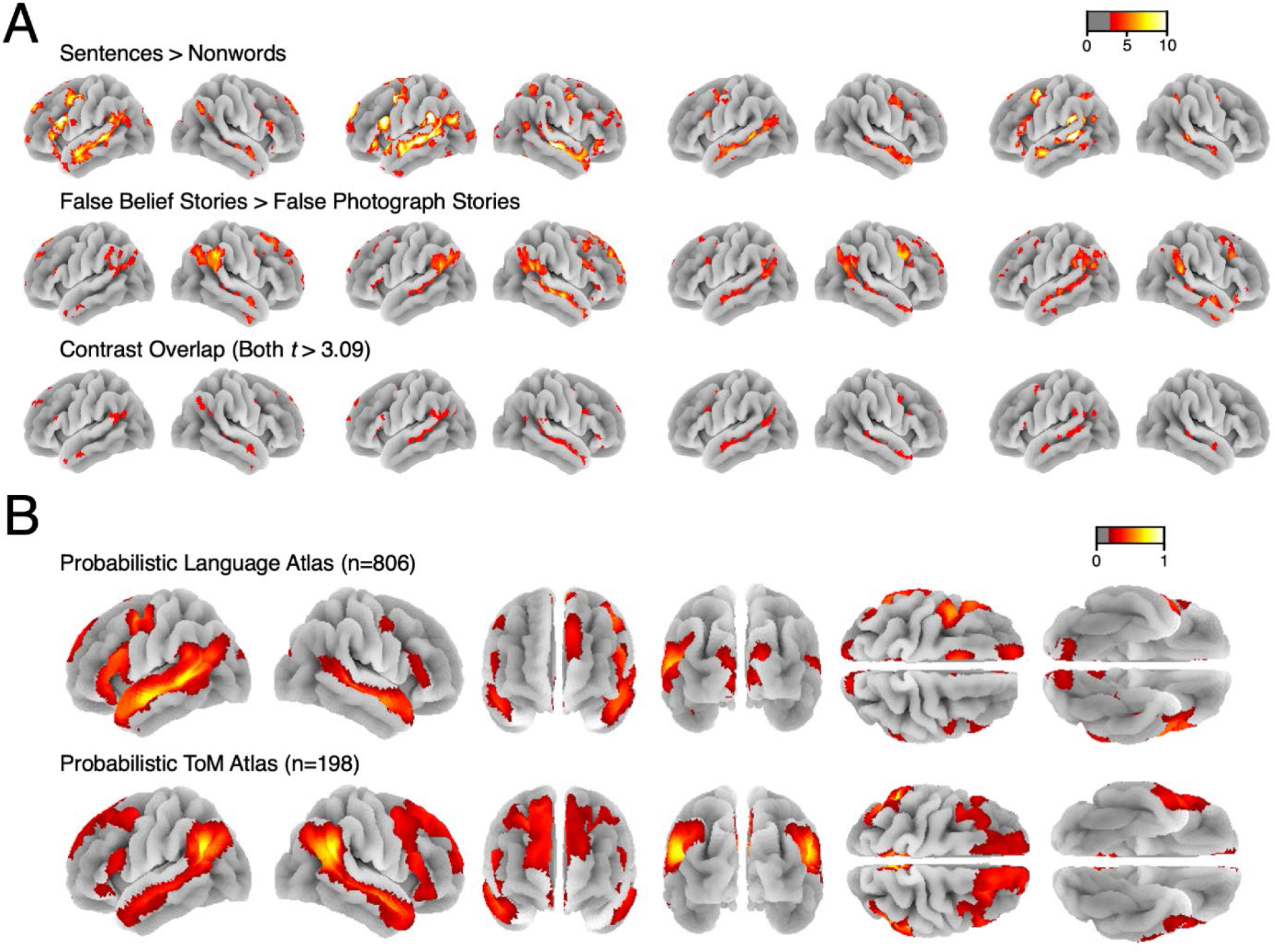
Comparison of whole-brain activation patterns for language (sentences > non-words) and ToM (false belief > false photo) contrasts. **A.** Responses to the language (top row) and ToM (middle row) localizer contrasts in four sample participants. Overlap (bottom row) is observed primarily in and around the angular gyrus/temporo-parietal junction (TPJ) and along the superior temporal sulcus, with some scattered overlap in the lateral frontal cortex. **B.** Whole-brain probabilistic atlases for the language and ToM localizer contrasts created from two large fMRI datasets by overlaying individual activation maps (see Lipkin et al., in press, for details). The two tasks elicit broadly similar spatial distributions of activity.

In an effort to clarify the relationship between the language and the ToM networks, we examine responses in the frontal and temporal language areas to the standard verbal ToM contrast (false belief stories > false photograph stories; Saxe & Kanwisher, 2003; the same contrast as was used in Deen et al., 2015), but also to a non-verbal ToM contrast (mental events > physical interactions in a rich naturalistic stimulus—a few-minute-long Pixar film; Jacoby et al., 2016). Jacoby et al. (2016) have previously shown that this non-verbal ToM contrast elicits a strong response in brain areas defined by the verbal ToM localizer (see also Richardson et al., 2018; Kamps et al., 2022).

In addition to our critical question about the involvement of core language network areas in ToM processing, we conduct exploratory analyses to investigate possible ToM responses in brain areas in the “periphery” of the language network—areas that show some response during language processing, but are less strongly integrated with the core left-hemisphere (LH) language areas, than those core areas are with one another (Fedorenko & Thompson-Schill, 2014; Chai et al., 2016). These peripheral regions include the right-hemisphere (RH) homotopes of the frontal and temporal LH areas and bilateral areas in the angular gyrus, and have been shown to differ in their functional profiles from the core language areas (e.g., Blank et al., 2016; Fedorenko et al., 2020; Ivanova et al., 2020 *inter alia*) and to show reduced static and dynamic functional correlations with the core language areas (e.g., Blank et al., 2014; Chai et al., 2016; Paunov et al., 2019; Braga et al., 2020). Further, these broad anatomical areas have been implicated by prior work in ToM, social processing, or social/affective aspects of language processing: the RH lateral frontal and temporal areas (e.g., Kaplan et al., 1990; Winner et al., 1998; Mitchell & Crow, 2005; Rajimehr et al., 2021; Hauptman et al., 2022), as well as bilateral angular gyri (e.g., Saxe & Kanwisher, 2003; Saxe, 2006, 2010; Lombardo et al., 2011; Mar, 2011; Schurz et al., 2014, 2017). These peripheral language areas may therefore show greater functional overlap with ToM reasoning, and thus possibly serve as transitional zones between the language-selective and the ToM-selective networks.

To foreshadow our results, we do not find that the core LH language areas support ToM reasoning. Although, similar to Deen et al. (2015), we find that the language areas respond to the verbal ToM contrast, we show i) that this effect is at least in part due to linguistic differences between the two conditions, and ii) that a non-verbal ToM condition does not engage the language network. In the language periphery, we find that non-verbal ToM elicits a strong response in both the RH homotopes of the language areas and in bilateral language-responsive areas in the angular gyrus. However, the detailed response profile of these areas differs from that of the ToM areas. Unlike the ToM areas, these peripheral language areas respond at least as strongly to depictions of social interactions with no mental state content as they do to depictions that encourage mentalizing. These results are therefore consistent with a broadly social function for peripheral language areas but do not support a role in ToM specifically.

## Materials & Methods

### General Approach

Our research design is informed by extensive prior evidence that the language and ToM networks are dissociable functional units in the human brain that can be reliably identified using a range of methods. *First*, a range of materials, tasks, and presentation formats yield remarkably stable definitions of both the language network (e.g., Fedorenko et al., 2010; Scott et al., 2017; Malik-Moraleda, Ayyash et al., in press) and the ToM network (Fletcher et al., 1995; Gallagher et al., 2000; Castelli et al., 2002; Sommer et al., 2007; Mason & Just, 2011; Jacoby et al., 2016, see e.g., Koster-Hale & Saxe, 2013, for review). Further, these networks emerge from task-free (resting state) functional correlation data (Braga & Buckner, 2017; Braga et al., 2020; DiNicola et al., 2020). These two networks generally show little spatial overlap, with areas of the superior temporal sulcus (STS) being the principal exception (Deen et al., 2015; Paunov et al., 2019). *Second*, the language and ToM networks show high within-network synchrony and lower between-network synchrony during both resting state and naturalistic language comprehension tasks (Paunov et al., 2019, 2022; Braga et al., 2020), supporting a functional dissociation between them. *Third*, language and ToM abilities show dissociable patterns of impairment: damage to the language network can impair language processing without impairing ToM reasoning (e.g., Dronkers et al., 1998; Varley et al., 2001; Apperly et al., 2006; Willems et al., 2011), whereas damage to the ToM network can impair ToM reasoning but without impairing language processing (e.g., Apperly et al., 2004; Martín-Rodríguez & León-Carrión, 2010; Domínguez et al., 2019), and language can be preserved in individuals whose ToM reasoning is otherwise impaired, as in some cases of autism spectrum disorders (e.g., Tager-Flusberg et al., 2005; Diehl et al., 2006) or schizophrenia (Sprong et al., 2007).

Based on the foregoing evidence, in this work, we assume the existence and (at least partial) functional dissociation of the language and theory of mind networks. We simply use “localizer task” contrasts (described below) as an efficient method to identify these networks in individual brains, in order to ask whether effects of interest are present (albeit to a lesser extent) in a given network (e.g., whether the language network shows evidence of mentalizing).

## Experimental Design

### Participants

162 individuals from the Cambridge/Boston, MA community participated for payment. All participants completed the language localizer (Fedorenko et al., 2010). They also all completed a verbal Theory of Mind (ToM) localizer task (Saxe & Kanwisher, 2003; task details are below, in **Materials and Procedure**). 160 of the 162 participants completed two runs of the verbal ToM task, and the remaining 2 participants completed a single run of the verbal ToM task. For the 160 participants who performed two runs, we evaluated the quality of the verbal ToM task data by examining the stability of the activation landscape across runs. In particular, we computed an across-runs spatial correlation across voxels that fall within the set of ToM masks corresponding to broad areas within which most participants show ToM responses (as described below in **Materials and Procedure**). Based on this analysis, we excluded 11 participants with negative spatial correlation values, leaving 149 participants. For the 2 participants who performed a single run of the verbal ToM task, we evaluated data quality by visual examination of the whole-brain activation maps for the localizer contrast (i.e., false belief > false photo; task details below); both participants’ maps looked as expected. Thus, overall, we include 151 participants in the analyses reported here (age 18-48, mean 24.7; 99 (66%) female). A subset of these participants (n=48) additionally completed a non-verbal ToM localizer task (Jacoby et al., 2016; task details below). (Of these, 34 participants completed both ToM localizer tasks within the same scanning session, whereas the remaining 14 participants completed them in different sessions.) Because at least two runs of a task are necessary to estimate the response magnitudes to the conditions of that task (to ensure independence between the data used to define the regions of interest and the data used to estimate the responses, as described below in **Materials and Procedure**), the two participants with a single run of the verbal ToM task were not included in the analyses of the verbal ToM task but they could still be used for defining the ToM fROIs and examining the responses in those fROIs to the conditions of the non-verbal ToM task.

138 of the 151 participants (∼91%) were right-handed, as determined by the Edinburgh handedness inventory (Oldfield, 1971), or self-report; the remaining participants (10 left-handed and 3 ambidextrous) showed typical left-lateralized language activations in the language localizer task (see Willems et al., 2014, for arguments for including left-handers in cognitive neuroscience experiments). 135 participants (∼89%) were native English speakers, the remaining 16 were fluent in English (see Malik-Moraleda, Ayyash et al., 2022 for evidence that language responses are similar in native and fluent speakers of English). All participants gave written informed consent in accordance with the requirements of MIT’s Committee on the Use of Humans as Experimental Subjects (COUHES).

### Materials and Procedure

For all participants, the scanning sessions lasted approximately two hours and included some tasks not related to the results reported here.

#### Language localizer

This task is described in detail in Fedorenko et al (2010) and subsequent studies from the Fedorenko lab (e.g., Fedorenko et al., 2011, 2020; Blank et al., 2014, 2016; Pritchett et al., 2018; Paunov et al., 2019; Shain, Blank, et al., 2020, among others; available for download from https://evlab.mit.edu/funcloc/). The language localizer targets higher-level aspects of language, including lexical and phrasal semantics, morphosyntax, and sentence-level pragmatic processing, to the exclusion of perceptual (speech or reading-related) and articulatory processes (see Fedorenko & Thompson-Schill, 2014, for discussion). Briefly, participants read sentences and lists of unconnected, pronounceable nonwords in a blocked design with a counterbalanced order across runs. Stimuli were presented one word/nonword at a time at the rate of 450ms per word/nonword. Participants read the materials passively and performed a simple button-press task at the end of each trial (included in order to help participants remain alert). Each block consisted of 3 6-s trials for total block duration of 18 sec. Each run consisted of 8 blocks per condition, with 5 14s rest periods (one at the beginning of the run, one at the end, and three interleaved between blocks). This localizer task has been extensively validated and shown to be robust to variation in the materials, modality of presentation, language, and task (Fedorenko et al., 2010; Scott et al., 2017; Malik-Moraleda et al., 2022; Ivanova et al., in prep.). Each participant completed two 5m 58s runs.

#### ToM localizer (verbal)

This task is described in detail in Saxe & Kanwisher (2003) and subsequent studies from the Saxe lab (e.g., Saxe & Wexler, 2005; Young et al., 2010; Bruneau, Pluta, et al., 2012, among others; available for download from http://saxelab.mit.edu/use-our-efficient-false-belief-localizer). The verbal ToM localizer targets “representational ToM” (Saxe, 2006), akin to “cognitive ToM” (Shamay-Tsoory et al., 2010; Dennis et al., 2013), that is, inferences about the propositional content of agents’ beliefs, desires, etc., to the exclusion of “affective ToM”, roughly, the capacity to understand and empathize with others’ emotional states (e.g., Brothers & Ring, 1992; Hein & Singer, 2008; Singer & Lamm, 2009). The task is based on the classic false belief paradigm (Wimmer & Perner, 1983; Wellman et al., 2001) and contrasts verbal vignettes about false beliefs (e.g., a protagonist has a false belief about an object’s location; the critical condition) vs. linguistically similar vignettes about false photo states (physical representations depicting outdated scenes, e.g., a photograph showing an object that has since been removed; the control condition). Participants read these vignettes, one at a time, in a slow event-related design. Each vignette was followed by a true / false comprehension question. This localizer task has been extensively validated and shown to be robust to variation in the materials, modality of presentation, and task (Saxe & Kanwisher, 2003; Saxe & Wexler, 2005; Saxe et al., 2006; Saxe & Powell, 2006; Young et al., 2010; Dodell-Feder et al., 2011; Bruneau, Dufour, et al., 2012; Koster-Hale & Saxe, 2013). 149 participants completed two runs, each lasting 4m 22s and consisting of 5 vignettes per condition. The remaining two participants who completed one run were excluded from the analysis of verbal ToM localizer activations, but were used for the analysis of responses of the ToM areas to the conditions of the non-verbal ToM localizer.

#### ToM localizer (non-verbal)

This non-verbal paradigm, based on a silent animated film, is described in detail in Jacoby et al. (2016) and subsequent studies (e.g., Richardson et al., 2018, 2020; Paunov et al., 2019, 2022). Similarly to the verbal ToM localizer, it targets brain regions that support inferences about others’ mental states, but in contrast to the main localizer, it is non-verbal, relying on participants engaging in mental state attribution from observed intentional actions. The task consists of passive viewing of an animated short film, Partly Cloudy (Pixar Animation Studios), which contains sections that are likely to elicit mental state attribution—the “mental” condition (e.g., a character falsely believes they have been abandoned by a companion)—as well as sections which simply depict physical events—the “physical” condition (e.g., a flock of storks flying). As described in Jacoby et al. (2016), such sections were identified and coded, by 5 independent coders, as the “mental” condition or the “physical” condition; two additional types of sections were identified and coded as “social” (i.e. characters interacting without strong mental or emotional dimensions, e.g., a cloud and stork playing) and “pain” (i.e. characters experiencing physical pain, e.g., a stork bitten by a crocodile). Jacoby et al. (2016) compared the activation patterns for the mental > pain contrast to those elicited by the verbal ToM contrast described above and found them to be similar in individual subjects. Here we primarily focus on the mental > physical contrast, which is conceptually more similar to the verbal ToM localizer contrast (Paunov et al., 2019). This localizer consists of a single run lasting 5m 48s, including 4 mental events with total duration 44s and 3 physical events with total duration 24s. Participants watched the film passively. (The localizer is available at http://saxelab.mit.edu/theory-mind-and-pain-matrix-localizer-movie-viewing-experiment; the Partly Cloudy short film itself must be purchased from Pixar Animation Studios.)

### fMRI data acquisition, preprocessing, and first-level modeling

#### Data acquisition

Whole-brain structural and functional data were collected on a whole-body 3 Tesla Siemens Trio scanner with a 32-channel head coil at the Athinoula A. Martinos Imaging Center at the McGovern Institute for Brain Research at MIT. T1-weighted structural images were collected in 176 axial slices with 1 mm isotropic voxels (repetition time (TR) = 2,530 ms; echo time (TE) = 3.48 ms). Functional, blood oxygenation level-dependent (BOLD) data were acquired using an EPI sequence with a 90° flip angle and using GRAPPA with an acceleration factor of 2; the following parameters were used: thirty-one 4.4 mm thick near-axial slices acquired in an interleaved order (with 10% distance factor), with an in-plane resolution of 2.1 mm × 2.1 mm, FoV in the phase encoding (A >> P) direction 200 mm and matrix size 96 × 96 voxels, TR = 2000 ms and TE = 30 ms. The first 10 s of each run were excluded to allow for steady state magnetization.

#### Preprocessing

fMRI data were analyzed using SPM12 (release 7487), CONN EvLab module (release 19b) and other custom MATLAB scripts. Each participant’s functional and structural data were converted from DICOM to NIFTI format. All functional scans were coregistered and resampled using B-spline interpolation to the first scan of the first session (Friston et al., 1995). Potential outlier scans were identified from the resulting subject-motion estimates as well as from BOLD signal indicators using default thresholds in CONN preprocessing pipeline (5 standard deviations above the mean in global BOLD signal change, or framewise displacement values above 0.9 mm, Nieto-Castanon, 2020). Functional and structural data were independently normalized into a common space (the Montreal Neurological Institute [MNI] template; IXI549Space) using SPM12 unified segmentation and normalization procedure (Ashburner & Friston, 2005) with a reference functional image computed as the mean functional data after realignment across all timepoints omitting outlier scans. The output data were resampled to a common bounding box between MNI-space coordinates (-90, -126, -72) and (90, 90, 108), using 2mm isotropic voxels and 4th order spline interpolation for the functional data, and 1mm isotropic voxels and trilinear interpolation for the structural data. Last, the functional data were then smoothed spatially using spatial convolution with a 4 mm FWHM Gaussian kernel.

#### First-level modeling

For both the language localizer task and the ToM localizer tasks, effects were estimated using a General Linear Model (GLM) in which each experimental condition was modeled with a boxcar function convolved with the canonical hemodynamic response function (HRF) (fixation was modeled implicitly, such that all timepoints that did not correspond to one of the conditions were assumed to correspond to a fixation period). Temporal autocorrelations in the BOLD signal timeseries were accounted for by a combination of high-pass filtering with a 128 seconds cutoff, and whitening using an AR(0.2) model (first-order autoregressive model linearized around the coefficient a=0.2) to approximate the observed covariance of the functional data in the context of Restricted Maximum Likelihood estimation (ReML). In addition to main condition effects, other model parameters in the GLM design included first-order temporal derivatives for each condition (included to model variability in the HRF delays), as well as nuisance regressors controlling for the effect of slow linear drifts, subject-motion parameters, and potential outlier scans on the BOLD signal.

### Definition of the language and ToM functional regions of interest

For each localizer, we defined a set of functional regions of interest (fROIs) using group-constrained, subject-specific localization (Fedorenko et al., 2010). For the core language fROIs, each individual map for the *sentences > nonwords* contrast was intersected with a set of five binary masks. These masks (available at OSF: https://osf.io/bzwm8/) were derived from a probabilistic activation overlap map for the same contrast in a large independent set of participants (n=220) using watershed parcellation, as described in Fedorenko et al. (2010) for a smaller set of participants. These masks included three in the left frontal cortex—in the inferior and middle frontal gyri—and two in the left temporal cortex (**Figure 3**). Within each mask, a participant-specific language fROI was defined as the top 10% of voxels with the highest *t*-values for the localizer contrast (see Lipkin et al., in press for evidence that the fROIs defined in this way are similar to the fROIs defined based on a fixed statistical significance threshold). In addition, we defined peripheral language fROIs: one in the left angular gyrus (using a mask derived in the same way as the masks for the core language areas) and six in the right hemisphere. Following e.g., Paunov et al. (2019), we defined masks for RH homotopes of core language areas and the RH AngG fROI by mirror-projecting the LH masks onto the right hemisphere and selecting the top 10% of most localizer-responsive voxels within these. In this way, the hemispheric symmetry only applies at the level of the masks; the particular sets of voxels selected within the LH vs. RH masks were free to differ within these symmetrical masks.

**Figure 2:**
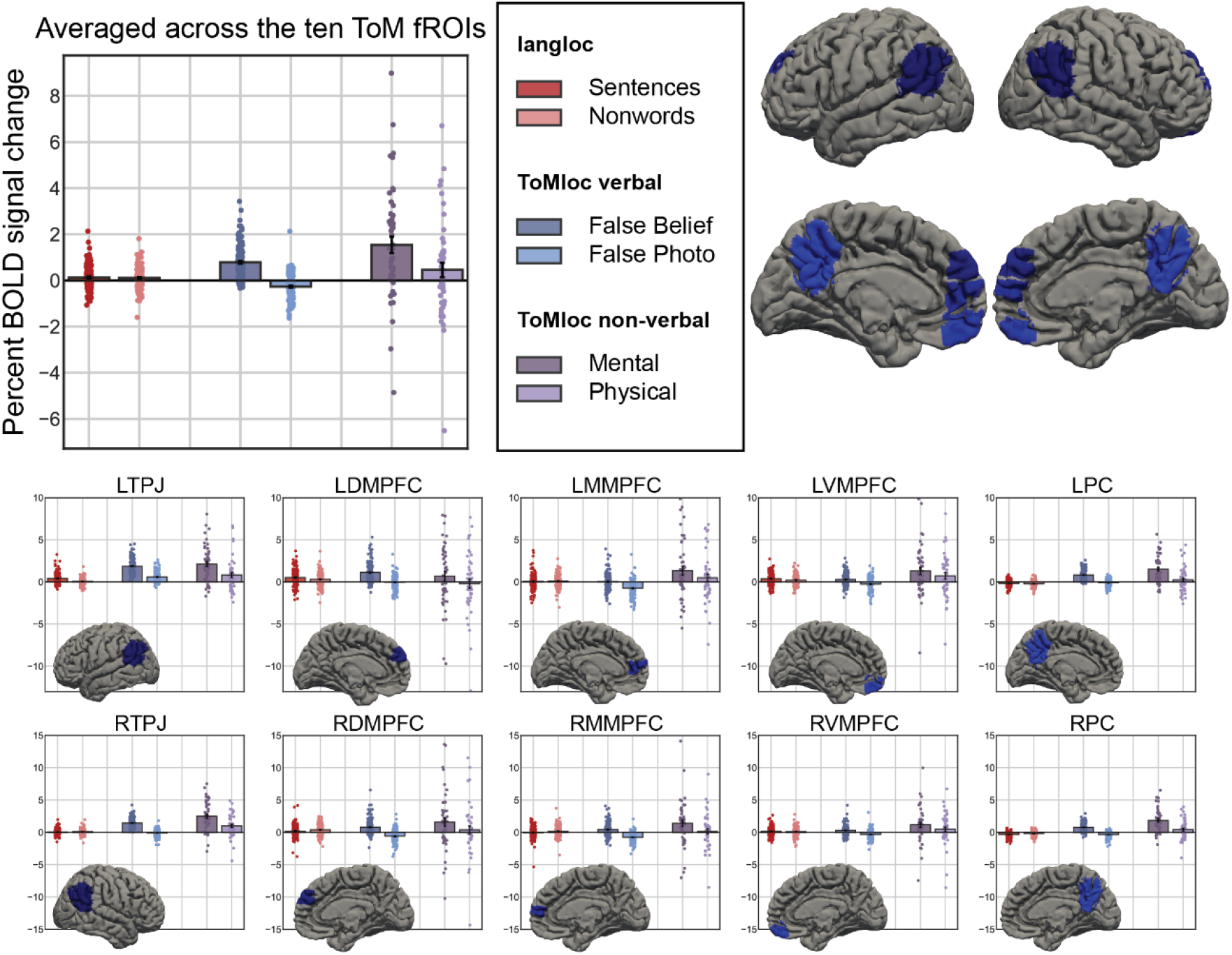
Responses to the conditions of the language localizer and the verbal and non-verbal ToM localizers in the theory of mind (ToM) network. The ToM network is not sensitive to the language contrast. ToM network activity increases in the presence of mental content, whether mediated verbally (false belief > false photo) or non-verbally (mental > physical). This overall pattern of results also holds within each of the 10 regions of the ToM network.

**Figure 3:**
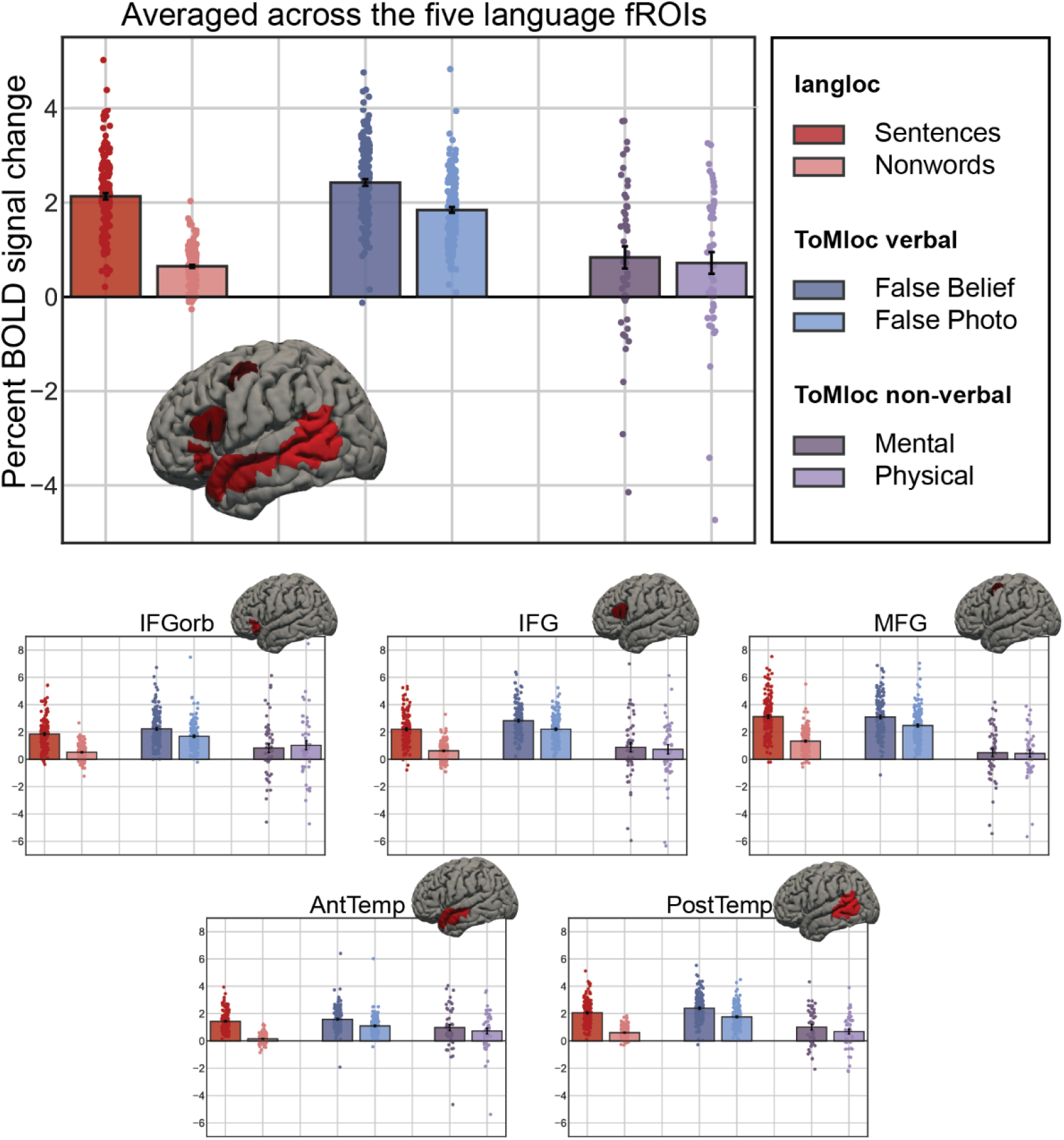
Responses to the conditions of the language localizer, and verbal and non-verbal ToM localizers in the language network. Replicating prior work, the language network shows a robust sentences > nonwords contrast and a weaker false belief > false photo contrast in the verbal ToM task. However, the language network shows no significant mental > physical contrast in the non-verbal ToM task, suggesting that the false belief > false photo contrast may be driven by linguistic differences between the two conditions. This overall pattern of results also holds within each of the 5 regions of the language network.

For the ToM fROIs, each individual map for the *false belief > false photo* contrast from the verbal ToM localizer was intersected with a set of ten binary masks (five in each hemisphere). These masks (available at OSF: https://osf.io/bzwm8/) were derived from a random effects map for the same contrast in a large independent set of 462 participants (Dufour et al., 2013). These masks included bilateral regions in temporoparietal junction, left and right precuneus / posterior cingulate cortex, and left and right dorsal, middle, and ventral medial prefrontal cortex (**Figure 2**). (Note that we did not include the masks for right superior temporal sulcus because (a) these areas are typically not included as part of the ToM network, though see Jacoby et al. (2016) for evidence that these areas respond to both verbal and nonverbal ToM contrasts, and (b) this mask largely subsumes the two temporal masks in the language network periphery). Within each mask, a participant-specific ToM fROI was defined as the top 10% of voxels with the highest *t*-values for the localizer contrast.

### Validation of the language and ToM fROIs

To ensure that the language and ToM fROIs behave as expected (i.e., language fROIs show a reliably greater response to the sentences condition compared to the nonwords condition; ToM fROIs show a reliably greater response to the false belief condition than the false photo condition), we used an across-runs cross-validation procedure (e.g., Nieto-Castañón & Fedorenko, 2012). In this analysis, the first run of the localizer was used to define the fROIs, and the second run to estimate the responses to the localizer conditions (in percent BOLD signal change, PSC), ensuring independence (e.g., Kriegeskorte et al., 2009); then the second run was used to define the fROIs, and the first run to estimate the responses; finally, the extracted magnitudes were averaged across the two runs to derive a single response magnitude for each of the localizer conditions. Statistical analyses were performed on these extracted PSC values.

## Statistical Analysis

We use PSC values derived from the localizer tasks to define dependent variables in linear mixed effects models in *lme4* (Bates et al., 2015) when examining entire networks, with random effects for *Participant* and *fROI,* or simple linear models when examining the fROIs separately. When examining the fROIs separately, reported *p*-values are adjusted for false discovery rate (Benjamini & Yekutieli, 2001) over the number of fROIs in the network. When examining the periphery of the language network, we treat the RH homotopic areas (n=5) and bilateral AngG areas (n=2) separately, and correct for the number of fROIs within each set.

### Units of Analysis

We conduct fROI-based analyses of critical effects in the core left-hemisphere language network (comprised of the LIFGorb, LIFG, LMFG, LAntTemp, and LPostTemp fROIs) and the bilateral ToM network (comprised of bilateral TPJ, DMPFC, MMPFC, VMPFC, and PC fROIs). We additionally conduct parallel analyses at the level of each individual language and ToM fROI. Finally, in exploratory analyses, we analyzed two key components of the “periphery” of the language network (Chai et al., 2016). This periphery is comprised of the RH homotopes of the LH language fROIs, and fROIs in the bilateral angular gyrus (AngG). As noted above, these fROIs have been shown in past work to systematically differ in their functional profiles from the core language regions and to show reduced static and dynamic functional correlations with the core regions (e.g., Blank et al., 2014; Chai et al., 2016; Braga et al., 2020). Moreover, of special relevance to the current investigation, these areas have previously been implicated in social cognition. In particular, the RH homotopes of the language areas have been argued to support social processing (e.g., Rajimehr et al., 2021), and the bilateral angular gyrus (AngG) is one of the key areas where overlap between language and ToM contrasts has been previously reported (Deen et al., 2015).

Because only a subset of our participants (48/151) completed the non-verbal ToM task, analyses that involve non-verbal contrasts are restricted to those participants that completed all three tasks (language localizer, verbal ToM localizer, and non-verbal ToM localizer). Otherwise, analyses included the 149 subjects with two ToM localizer runs.

### Main Analyses

Our localizers provide the following key conditions:

- **Language localizer:** Sentences, Nonwords
- **Verbal ToM localizer:** False Belief, False Photo
- **Non-verbal ToM localizer:** Mental, Physical, Social, Pain

In language areas, sentences should elicit a larger response than nonwords. In ToM areas, false belief stories should elicit a larger response than false photo stories, and video segments with mental content should elicit a larger response than video segments with physical content (as well as segments that depict physical pain (Jacoby et al., 2016), and, to a lesser extent, segments that depict social interactions). Critically, if the language areas support some aspects of ToM, they should also show false belief > false photo and mental > physical effects.

To test a contrast, we use the following linear mixed effects model, where *PSC* reflects the percent BOLD signal change (relative to the fixation baseline) associated with a given condition in a specific participant and fROI, *Contrast* is a binary indicator variable indexing which condition of the critical contrast is reflected by the PSC (e.g., whether the PSC measures the response to nonwords or sentences in the sentences > nonwords contrast), and the critical bolded variable is evaluated with a likelihood ratio test:

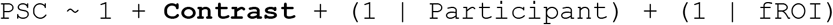

A fixed-effects only variant of this model is used for tests in individual fROIs:

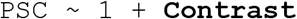

Note that we could have included by-participant and by-fROI random slopes for *Contrast* in the network model and a by-participant random intercept in the individual fROI model, but we found that doing so led to frequent problems with model identification in critical tests (non-convergence or singular fits).

### Linguistic Analyses

To better understand any linguistic determinants of verbal ToM effects in language regions, we analyzed the verbal ToM materials in terms of linguistic properties that are known, based on past behavioral and neural findings, to modulate language network activity. If the false belief conditions differ systematically from the false photo conditions in one or more of these dimensions, these differences could account for differences in language network activation. We considered the following linguistic predictors:

- **Num Words:** The number of words in an item. Language network activity has previously been associated with the length of linguistically coherent spans (e.g., Pallier et al., 2011; Fedorenko et al., 2016).
- **Num Sents:** The number of sentences in an item, which may modulate language network activity via sentence wrap-up processes (e.g., Just & Carpenter, 1980; Rayner et al., 2000).
- **Constituent End:** Whether a word terminates a syntactic constituent in a hand-corrected phrase structure tree. Constituent boundaries may modulate language network activity via constituent wrap-up processes (Nelson et al., 2017).
- **Integration Cost:** A measure of working memory retrieval difficulty. Integration cost is posited by the Dependency Locality Theory (Gibson, 2000) as an account of word-by-word variation in the difficulty of building linguistic representations in working memory. Here we use a variant of DLT integration cost that has been associated with language network activity in prior work (Shain, Blank, et al., in press).
- **Unigram Surprisal:** A measure of word frequency, specifically: the negative log of a word’s marginal probability according to a unigram KenLM language models (Heafield et al., 2013) trained on the Gigaword 3 corpus (Graff et al., 2007). Stronger language network activation has been associated with less frequent words (higher unigram surprisal, e.g. Schuster et al., 2016).
- **5-gram Surprisal:** A measure of word predictability, specifically: the negative log probability of a word in context according to a 5-gram KenLM language model (Heafield et al., 2013) trained on the Gigaword 3 corpus (Graff et al., 2007). Stronger language network activation has been associated with less predictable words (higher 5-gram surprisal, e.g. Lopopolo et al., 2017; Shain, Blank, et al., 2020).
- **PCFG Surprisal:** A measure of word predictability, specifically: the negative log probability of a word in context according to a probabilistic phrase structure grammar (PCFG) parser (van Schijndel et al., 2013) trained on a generalized categorical grammar reannotation (Nguyen et al., 2012) of the Wall Street Journal portion of the Penn Treebank corpus (Marcus et al., 1993). PCFG and 5-gram Surprisal effects have been shown to be dissociable in the human language network (Shain, Blank, et al., 2020).

Item-level values for Constituent End, Integration Cost, Unigram Surprisal, 5-gram Surprisal, and PCFG Surprisal were computed by averaging their respective values over all words in an item.

We ask whether controlling for these linguistic variables attenuates the false belief > false photo contrast in the language network. To investigate this question, we first regress each variable individually out of the item-wise PSCs in each language region of each participant. We then compute the change in the false belief > false photo contrast (i.e., the change in the difference between the average response to false belief items and the average response to false photo items) due to a linguistic control. For simplicity, we refer to the change due to linguistic feature *X* in the false belief > false photo effect as ΔToM.*X*. To test ΔToM.*X* for significance in a given functional network, we model it as the dependent variable in linear mixed effects models with the following structure, where the fixed intercept is evaluated with a likelihood ratio test:

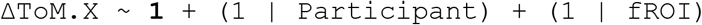

Similarly, we also examine the combined effect of regressing out all control variables simultaneously. In individual fROIs, we test ΔToM.*X* with a one-sample *t*-test.

## Results

### The Theory of Mind Network Shows Both Verbal and Non-verbal ToM Effects

Replicating prior work (e.g., Jacoby et al., 2016), our results show that functionally localized regions previously associated with ToM reasoning are significantly more activated in the presence of mental state content (**Figure 2)**, whether this content is delivered verbally (false belief > false photo) or nonverbally (mental > physical). The false belief > false photo contrast is significant in the network overall (β = 1.05, *p* < 0.001***) as well as in each individual ToM fROI (**Table 1**). The mental > physical contrast is significant overall (β = 1.09, *p* < 0.001***) and numerically positive in each individual ToM fROI, achieving significance in the TPJ and PC fROIs bilaterally (**Table 1**). The response of the ToM network to segments with mental state content is significantly larger than to segments that depict non-mental forms of social interaction (mental > social: β = 0.40, *p* = 0.001*; **Figure 8**) or to segments in which characters experience physical pain (i.e., “affective ToM”, mental > pain: β = 1.15, *p* < 0.001***), supporting a selective role for this network in the mentalizing aspects of ToM (“cognitive ToM”; Saxe & Powell, 2006; Bruneau, Pluta, et al., 2012). Note that we did not evaluate the language localizer contrast in the ToM network statistically because our localizer materials are not controlled for mental state content, but as can be seen in Figure 2, responses to these conditions are generally low (see also Koster-Hale & Saxe, 2013 and Deen et al., 2015, who show the lack of engagement of the ToM network for sentences devoid of mental/social content, in the presence of robust responses to those stimuli in the language network).

**Table 1:**
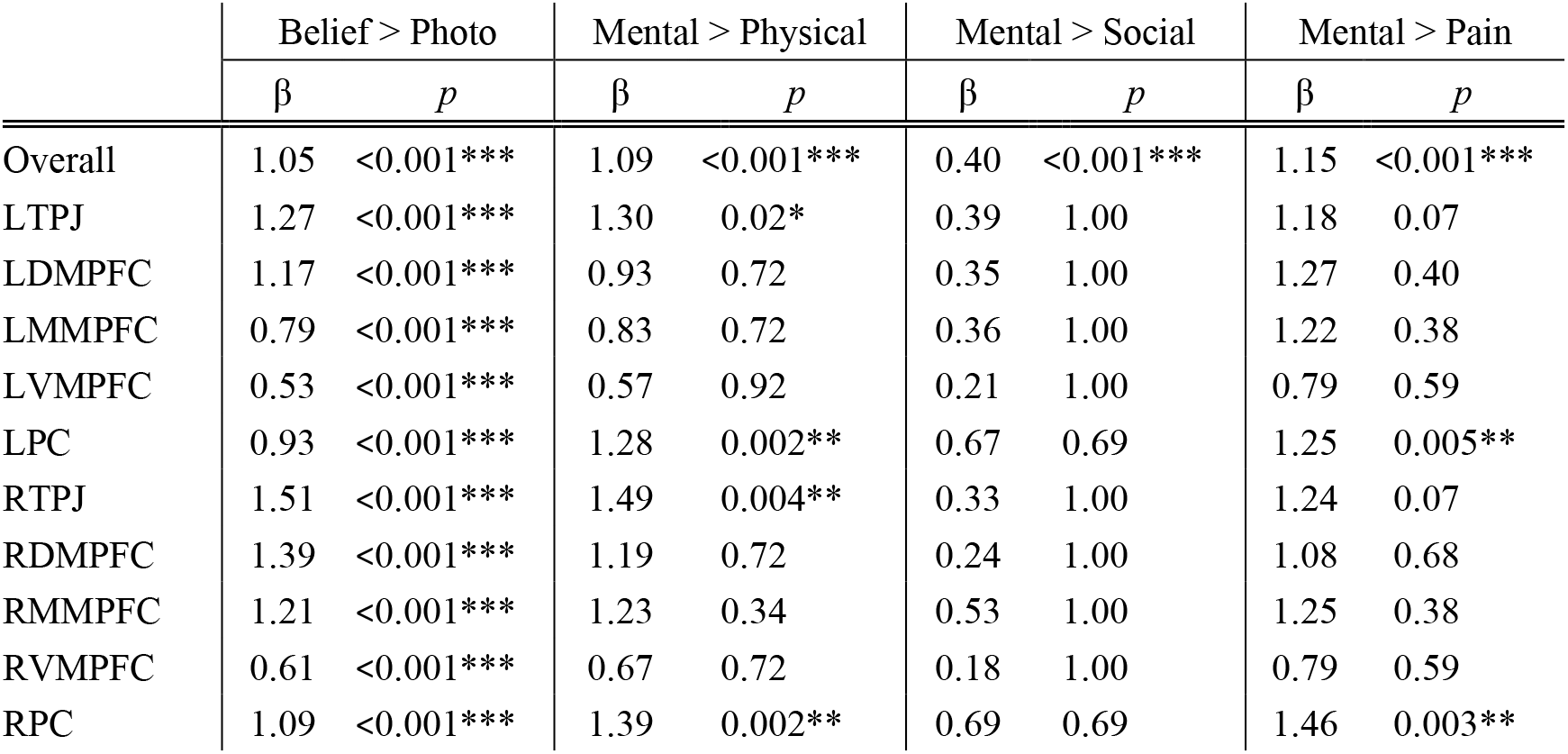
Size and significance of key contrasts in the ToM network (Overall) and each of its 10 component fROIs (fROI-level *p* values are FDR-corrected for 10 fROIs). Numerical estimates (β) and network-wide significance tests show a selective response to mentalizing across verbal and non-verbal representation formats.

### The Language Network Shows Verbal but not Non-verbal ToM Effects

Replicating much prior work, the core left-hemisphere language network (**Figure 3)** shows a larger response to sentences over nonword lists (Fedorenko et al., 2010), and replicating Deen et al. (2015), to false belief items over false photo items in the verbal ToM localizer. Both of these contrasts are significant in the language network as a whole (sentences > nonwords: β = 1.49, *p* < 0.001***; false belief > false photo: β = 0.58, *p* < 0.001***) and in each individual language fROI (**Table 2**). The overall response to both conditions of the verbal ToM localizer is more similar to the response to sentences than to the response to nonwords (both ToM conditions are of course presented in coherent language), and indeed both verbal ToM conditions elicit a significantly larger response than the nonwords condition of the language localizer, both in the language network as a whole (false belief > nonwords: β = 1.78, *p* < 0.001***; false photo > nonwords: β = 1.20, *p* < 0.001***) and in each individual language fROI.

**Table 2:** Size and significance of key contrasts in the language network (Overall) and each of its 5 component fROIs (fROI-level *p* values are FDR-corrected). Numerical estimates (β) and network-wide significance tests show a selective response to language (Sentence > Non-word) and verbal ToM (Belief > Photo), but no robust response to non-verbal ToM (Mental > Physical) or selectivity for mentalizing over social interactions (Mental < Social).

|  | Sent > Nonwd |  | Belief > Photo |  | Mental > Physical |  | Mental > Social |  | Mental > Pain |  |
| --- | --- | --- | --- | --- | --- | --- | --- | --- | --- | --- |
| | $\beta$ | $p$ | $\beta$ | $p$ | $\beta$ | $p$ | $\beta$ | $p$ | $\beta$ | $p$ |
| Overall | 1.49 | <0.001*** | 0.58 | <0.001*** | 0.12 | 0.29 | -0.25 | 0.03* | 0.29 | 0.009** |
| LIFGop | 1.34 | <0.001*** | 0.54 | <0.001*** | -0.20 | 1.00 | -0.36 | 1.00 | 0.45 | 1.00 |
| LIFG | 1.58 | <0.001*** | 0.62 | <0.001*** | 0.15 | 1.00 | -0.28 | 1.00 | 0.16 | 1.00 |
| LMFG | 1.78 | <0.001*** | 0.62 | <0.001*** | 0.05 | 1.00 | -0.43 | 1.00 | -0.05 | 1.00 |
| LAntTemp | 1.27 | <0.001*** | 0.49 | <0.001*** | 0.25 | 1.00 | 0.00 | 1.00 | 0.44 | 0.98 |
| LPostTemp | 1.45 | <0.001*** | 0.63 | <0.001*** | 0.33 | 1.00 | -0.18 | 1.00 | 0.47 | 0.98 |

However, the ToM effect is greatly attenuated when using a non-verbal (mental > physical) contrast. Neither the language network as a whole (mental > physical: β = 0.12, *p* = 0.29) nor any fROI within it registers a significant effect. Furthermore, the language network is significantly less responsive to segments with mental content than to segments that depict non-mental social interactions (mental > social: β = -0.25, *p* = 0.03*; **Table 2**). Thus, unlike the ToM network, the language network is not selectively engaged by reasoning about the content of others’ minds relative to other kinds of social cognition. A notable exception to the pattern of non-selectivity for mentalizing is the fact that the language network is significantly more responsive to segments that depict mental content than to segments that involve physical pain (mental > pain: β = 0.29, *p* = 0.009**). We revisit this finding in the discussion.

In addition, the overall response to the conditions of the non-verbal ToM localizer is more similar in magnitude to the response to nonwords than to the response to sentences, and indeed both non-verbal ToM conditions elicit a significantly smaller response than the sentences condition of the language localizer, both in the language network as a whole (sentences > mental: β = 1.33, *p* < 0.001***; sentences > physical: β = 1.44, *p* < 0.001***) and in most individual language fROIs (**Table 3**).

**Table 3:** Size and significance of contrasts between sentences and mental/physical conditions of the non-verbal ToM localizer in the language network (Overall) and each of its 5 component fROIs (fROI-level *p* values are FDR-corrected). Responses to sentences are significantly larger in the language network as a whole (Overall) and in most of its 5 component fROIs.

|  | Sent > Mental |  | Sent > Physical |  |
| --- | --- | --- | --- | --- |
| | $\beta$ | $p$ | $\beta$ | $p$ |
| Overall | 1.33 | <0.001*** | 1.44 | <0.001*** |
| LIFGop | 1.07 | 0.01* | 0.87 | 0.05 |
| LIFG | 1.24 | 0.005** | 1.39 | 0.002** |
| LMFG | 2.75 | <0.001*** | 2.80 | <0.001*** |
| LAntTemp | 0.49 | 0.11 | 0.74 | 0.006** |
| LPostTemp | 1.09 | <0.001*** | 1.42 | <0.001*** |

### Linguistic Features Explain Verbal ToM Effects in the Language Network

The fact that ToM effects only emerge in the language network for a verbal contrast (cf. the ToM network, where both verbal and non-verbal ToM contrasts elicit an effect) suggests that this effect may reflect *linguistic* differences between the false belief and false photo conditions of the verbal ToM localizer task, rather than theory of mind reasoning. If true, this hypothesis predicts i) that the false belief and false photo conditions will systematically differ in linguistic features that modulate language network activity independently of ToM, and thus ii) that controlling for the relevant features will attenuate the false belief > false photo effect in the language network. To test this hypothesis, we analyzed the effect of controlling for seven independently motivated linguistic features on the size of the false belief > false photo contrast: *Num Words*, *Num Sents*, and item-wise averages of *Constituent End*, *Integration Cost*, *Unigram Surprisal*, *5-gram Surprisal*, *Probabilistic Context-Free Grammar (PCFG) Surprisal*, for definitions, see **Linguistic Analyses**). The distributions of these features in the false belief and false photo materials are visualized in **Figure 4A**. As shown, false belief items are systematically higher in dimensions that are known to modulate language network activity, including *Num Words*\*, *Num Sents*\*, *Integration Cost*\*, *5-gram Surprisal*\*, *PCFG Surprisal* (stars indicate significant differences in a 2-sample *t*-test). These feature distributions are consistent with our hypothesis that the verbal ToM contrast is confounded with linguistic complexity.

**Figure 4:**
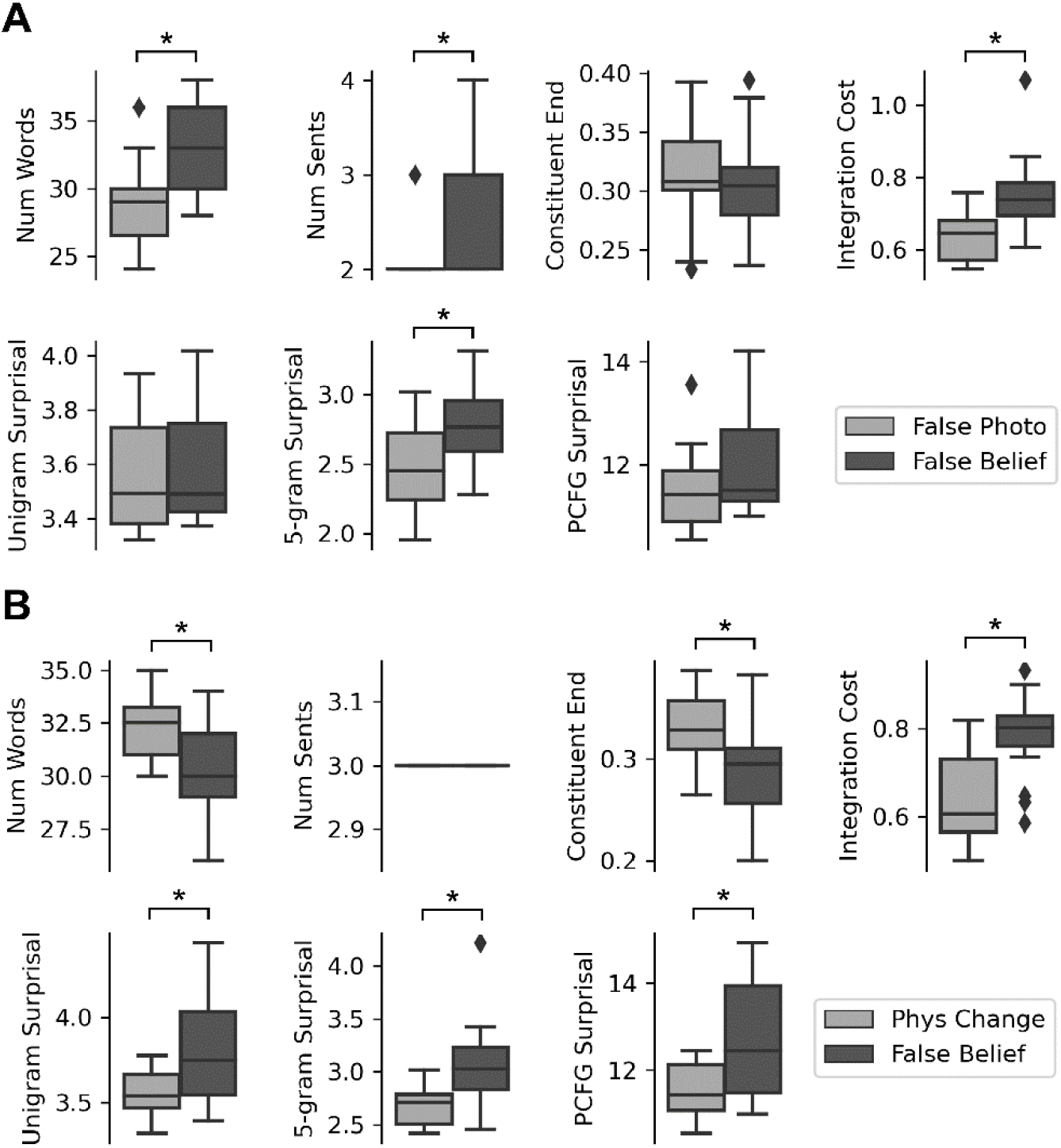
**A.** Distribution of linguistic features in the false belief vs. false photo items of the verbal ToM localizer task. The false belief items are systematically higher in dimensions that are known to modulate language network activity, including number of words, number of sentences, integration cost, and 5-gram surprisal. **B.** Identical analyses for the *false belief* vs. *physical change* items of Deen et al. (2015). These items also differ significantly along dimensions known to modulate language network activity, especially integration cost, unigram surprisal, 5-gram surprisal, and PCFG surprisal.

To test the hypothesis directly, we analyzed the impact of controlling for each linguistic feature individually, as well as the impact of controlling for all seven linguistic features jointly, on the magnitude of the verbal ToM contrast in the language network (for statistical procedures, see **Statistical Analysis**). Results are plotted in **Figure 5**. The predictors *Num Words*, *Num Sents*, *Constituent End*, *Integration Cost*, and *5-gram Surprisal* significantly attenuate the verbal ToM contrast in the language network as a whole as well as in each fROI within it (**Table 4**). *PCFG Surprisal* significantly attenuates the ToM contrast in the entire network, as well as in LMFG and LPostTemp. *Unigram Surprisal* does not have a significant effect on the ToM contrast (see Shain, 2019 for related findings). In addition, jointly controlling for all linguistic features attenuates the verbal ToM contrast by 0.49 (*p* < 0.001***) network-wide. Given that the network-wide false belief > false photo contrast is 0.58, this means that linguistic differences account for at least 84% of the verbal ToM effect in the language network, rendering its residualized effect size (0.09) comparable to that of the nonverbal mental > physical contrast in the language network (0.11). Although the attenuated verbal ToM effect remains significant in the left-hemisphere language network overall and in each component fROI (p < 0.001***), our analysis only considered a handful of linguistic variables and therefore only provides a lower bound on the proportion of verbal ToM contrast that is attributable to linguistic factors.

**Figure 5:**
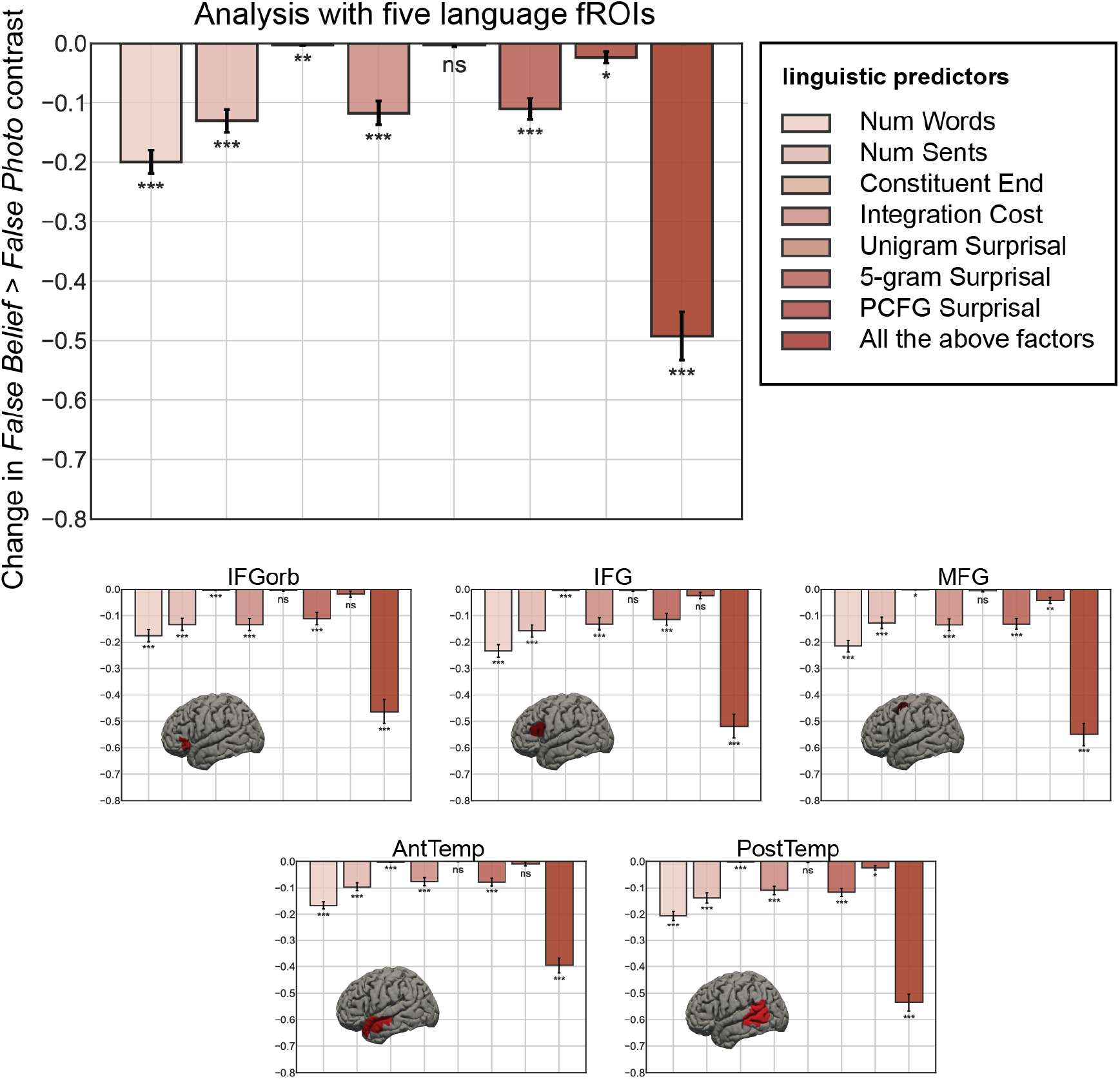
Effects of controlling for linguistic features on the false belief > false photo contrast in the language network. Effects are universally negative, meaning that controlling for the variable systematically attenuates verbal ToM contrasts throughout the language network.

**Table 4:**
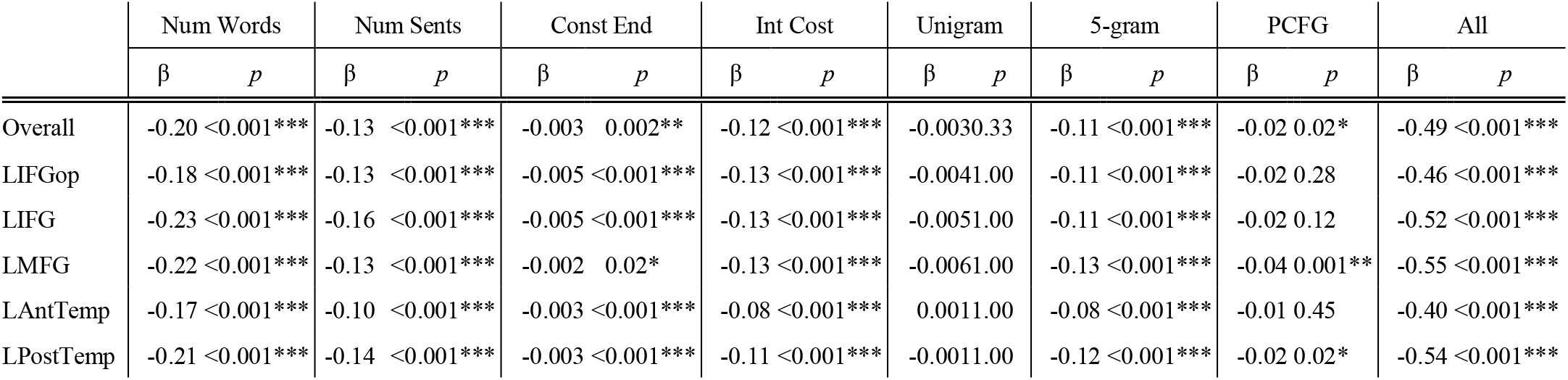
Effects of controlling for linguistic variables in the core language network (fROI-level *p* values are FDR-corrected). Effect estimates (β) represent the change in the verbal ToM contrast (false belief > false photo) due to controlling for a linguistic variable. Most variables we considered significantly attenuate the ToM contrast in the language network as a whole and in most of its 5 component fROIs.

Deen et al. (2015) also reported language-ToM overlap using a broader but more carefully linguistically controlled contrast between an additional set of stories describing false beliefs and a set of stories describing physical changes with no mental state attribution. Although this contrast cannot adjudicate between ToM and other forms of social cognition, its overlap with the language system suggests that at least some language-responsive areas may be recruited for some social cognitive functions, including possibly ToM. Deen et al. (2015) fixed the number of sentences in each item at 3 and controlled for a diverse set of linguistic features: “number of words, mean syllables per word, Flesch reading ease, number of noun phrases, number of modifiers, number of higher level constituents, number of words before the first verb, number of negations, and mean semantic frequency (log Celex frequency)” (Deen et al., 2015, p. 4598). Nonetheless, as shown in **Figure 4B**, the false belief items differ statistically from the physical change items along every relevant dimension in our linguistic evaluation (number of sentences has no variance by construction, as noted above), and the false belief items are systematically higher in dimensions known to increase language network activity: they have higher average integration cost, involve less frequent words (higher unigram surprisal), and are less predictable on the basis of both word co-occurrences (5-gram surprisal) and syntactic structure (PCFG surprisal). Thus, differences between these conditions in language-selective areas are also plausibly driven by linguistic confounds, despite considerable effort invested in matching along many linguistic features. The methodological upshot of this outcome is that linguistic matching of complex verbal stimuli is challenging, if not impossible, due to the myriad structural and statistical relationships that hold between words in language. For designs that seek to study modulation of language-selective brain areas by content-related (semantic) contrasts, it may be necessary to avoid verbal stimuli, or at least to supplement verbal contrasts with non-verbal ones.

### The Language Network’s “Periphery” May Support Broader Social Cognition

Even though the core LH language areas do not show evidence of supporting mental state attribution in our study, it has been argued that some regions in the periphery of the language processing system (Chai et al., 2016) are associated with ToM reasoning and/or social processing more generally. Here we consider two candidate components of the language periphery: the right-hemisphere homotopes of the LH core language regions and the language-responsive areas in the bilateral angular gyri. The function(s) of both of these components remains debated in the field (see **Discussion**).

Responses to the key conditions of all three localizers in the right hemisphere (RH) homotopes of the core language areas are plotted in **Figure 6**. RH language regions show considerably less selectivity for language processing than their LH counterparts: unlike in the LH, linguistic stimuli (the sentence condition of the language localizer, and the conditions of the verbal ToM localizer) elicit lower responses than the rich visual stimuli from the non-verbal ToM localizer (see Small, Lipkin et al., 2021 and Ivanova in prep.-b for additional evidence of lower selectivity of the RH language regions). In addition, unlike the core language network but similar to the ToM network, these RH language areas respond significantly to both ToM contrasts (false belief > false photo: β = 0.59, *p* < 0.001***; mental > physical: β = 0.38, *p* = 0.001**). However, unlike the ToM network, RH language areas are not selective for mentalizing segments relative to segments that depict physical pain (mental > pain: β = -0.01, *p* = 0.94), and they are significantly less responsive to mentalizing segments than to segments that depict non-mental forms of social interaction (mental > social: β = -0.28, *p* = 0.019*) (**Figure 6**, **Table 5**). Thus, any contribution of the RH language areas to social cognition is not restricted to cognitive ToM/mentalizing.

**Figure 6:**
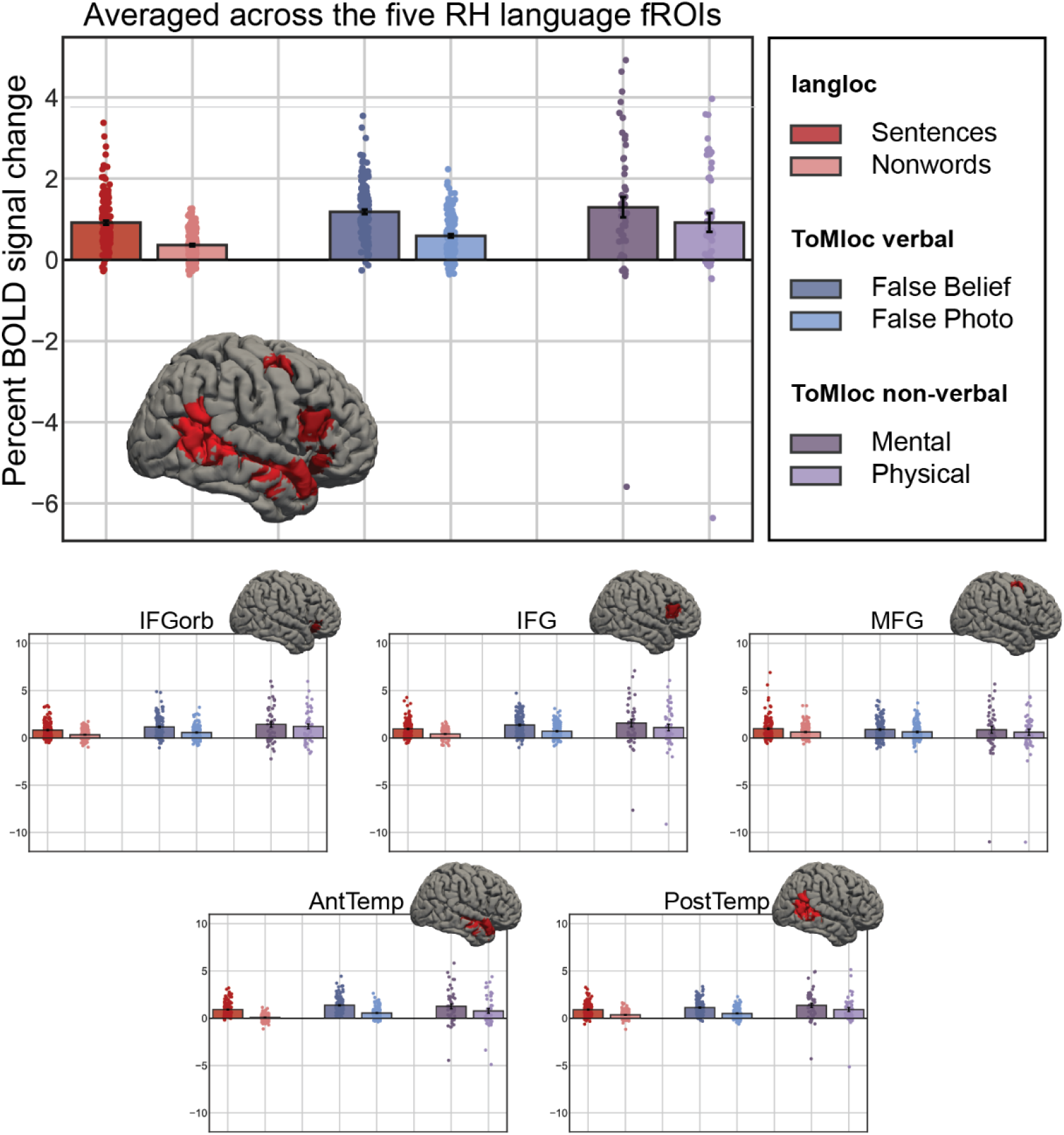
Responses to the conditions of the language localizer, and verbal and non-verbal ToM localizers in the RH homotopes of the language network. Replicating prior work, RH language regions show a robust sentences > nonwords contrast. However, unlike the core LH language network (Figure 3), RH language regions show similarly strong responses to both the false belief > false photo contrast of the verbal ToM localizer and the mental > physical contrast of the non-verbal ToM localizer. This overall pattern of results also holds within each of the 5 RH homotopes of the core LH language network.

**Table 5:** Size and significance of key contrasts in the language periphery, comprised of language-responsive fROIs in (i) the right hemisphere homotopes of core language areas and (ii) bilateral angular gyri (fROI-level *p* values are FDR-corrected). Like the language network, these areas respond to language (Sentence > Non-word), and like the ToM network, they respond to verbal (Belief > Photo) and non-verbal (Mental > Physical) ToM. However, unlike the ToM network, they do not respond more to mentalizing relative to other forms of social interaction (null or negative Mental > Social effects), and the RH homotopic areas furthermore do not respond more to mentalizing relative to observing physical pain (Mental > Pain). These patterns are not consistent with a selective response to ToM, but could be consistent with a more broadly social function in addition to language (but see Discussion for an alternative interpretation).

|  | Sent > Nonwd |  | Belief > Photo |  | Mental > Physical |  | Mental > Social |  | Mental > Pain |  |
| --- | --- | --- | --- | --- | --- | --- | --- | --- | --- | --- |
| | $\beta$ | $p$ | $\beta$ | $p$ | $\beta$ | $p$ | $\beta$ | $p$ | $\beta$ | $p$ |
| RH Overall | 0.55 | <0.001*** | 0.59 | <0.001*** | 0.38 | <0.001*** | -0.29 | 0.02* | -0.01 | 0.94 |
| RIFGop | 0.49 | <0.001*** | 0.60 | <0.001*** | 0.22 | 1.00 | -0.33 | 1.00 | -0.14 | 1.00 |
| RIFG | 0.53 | <0.001*** | 0.66 | <0.001*** | 0.47 | 1.00 | -0.34 | 1.00 | -0.15 | 1.00 |
| RMFG | 0.36 | 0.001** | 0.26 | 0.04* | 0.26 | 1.00 | -0.34 | 1.00 | -0.18 | 1.00 |
| RAntTemp | 0.85 | <0.001*** | 0.82 | <0.001*** | 0.48 | 1.00 | -0.14 | 1.00 | 0.25 | 1.00 |
| RPostTemp | 0.55 | <0.001*** | 0.62 | <0.001*** | 0.44 | 1.00 | -0.30 | 1.00 | 0.17 | 1.00 |
| AngG Overall | 0.46 | <0.001*** | 0.61 | <0.001*** | 0.73 | <0.001*** | 0.01 | 0.96 | 0.68 | <0.001*** |
| LAngG | 0.79 | <0.001*** | 0.75 | <0.001*** | 0.90 | 0.20 | 0.10 | 1.00 | 0.89 | 0.26 |
| RAngG | 0.14 | 0.03* | 0.47 | <0.001*** | 0.55 | 0.21 | -0.08 | 1.00 | 0.47 | 0.39 |

Responses to the conditions of all three localizers in the language-responsive areas in the angular gyrus (bilaterally) are plotted in **Figure 7**. Like the ToM network, these areas respond significantly to both ToM contrasts (false belief > false photo: β = 0.61, *p* < 0.001***; mental > physical: β = 0.73, *p* < 0.001***). In addition, the AngG language areas are selective for mentalizing segments relative to segments that depict physical pain (mental > pain: β = 0.68, *p* < 0.001***). However, unlike the ToM network, these areas do not show a mental > social effect (β = 0.01, *p* = 0.96) (**Figure 7**, **Table 5**). Thus, similar to what we observed for the RH language areas, any contribution of the language areas in the AngG to social cognition appears to be different from that of the ToM network in that it is not restricted to ToM reasoning.

**Figure 7:**
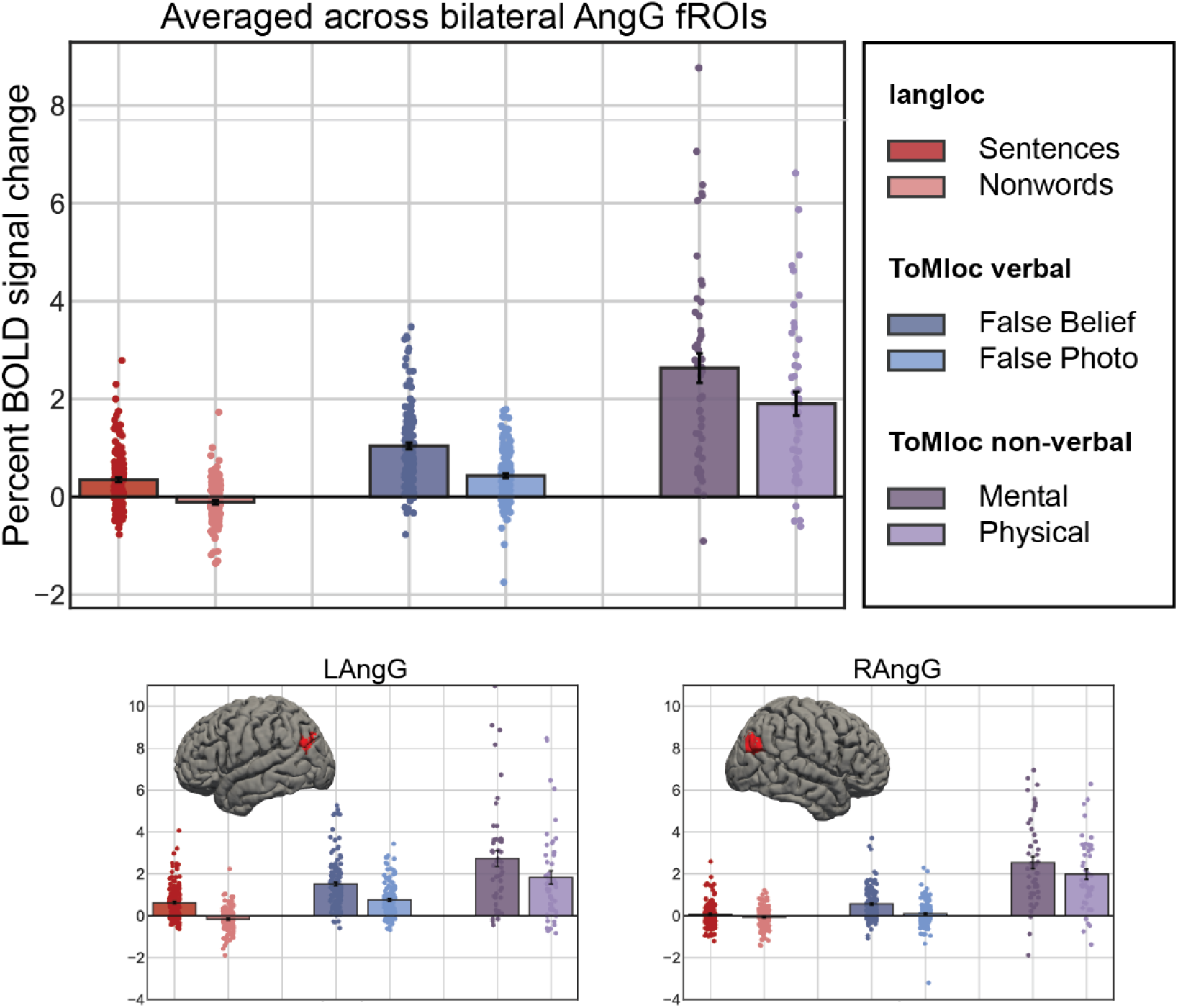
Responses to the conditions of the language localizer, and verbal and non-verbal ToM localizers in the bilateral angular gyri. Unlike the core LH language network (Figure 3), these regions show similarly strong responses to both the false belief > false photo contrast of the verbal ToM localizer and the mental > physical contrast of the non-verbal ToM localizer. This overall pattern of results holds in both the LH and RH AngG fROIs.

**Figure 8** shows responses in the four sets of fROIs examined here to the four conditions of the non-verbal ToM localizer (*mental*, i.e. segments depicting mental state content; *physical*, i.e. segments depicting physical events; *social*, i.e. segments depicting non-mentalizing social interactions; and *pain*, i.e. segments depicting physical pain). Only the ToM network shows the characteristic profile of greater response to the *mental* condition than either the *physical* or *social* conditions; in the language network and its periphery, the response to the *social* condition is at least as large as the response to the *mental* condition. The language network response to all conditions in the task is lower than that of the other networks, likely due to the fact that this task is entirely non-verbal. The RH language homotopes and the angular gyri both show a stronger response to the *mental* and *social* conditions than to the *physical* and *pain* conditions, consistent with a broadly social function for these areas, rather than a ToM-selective one.

**Figure 8:**
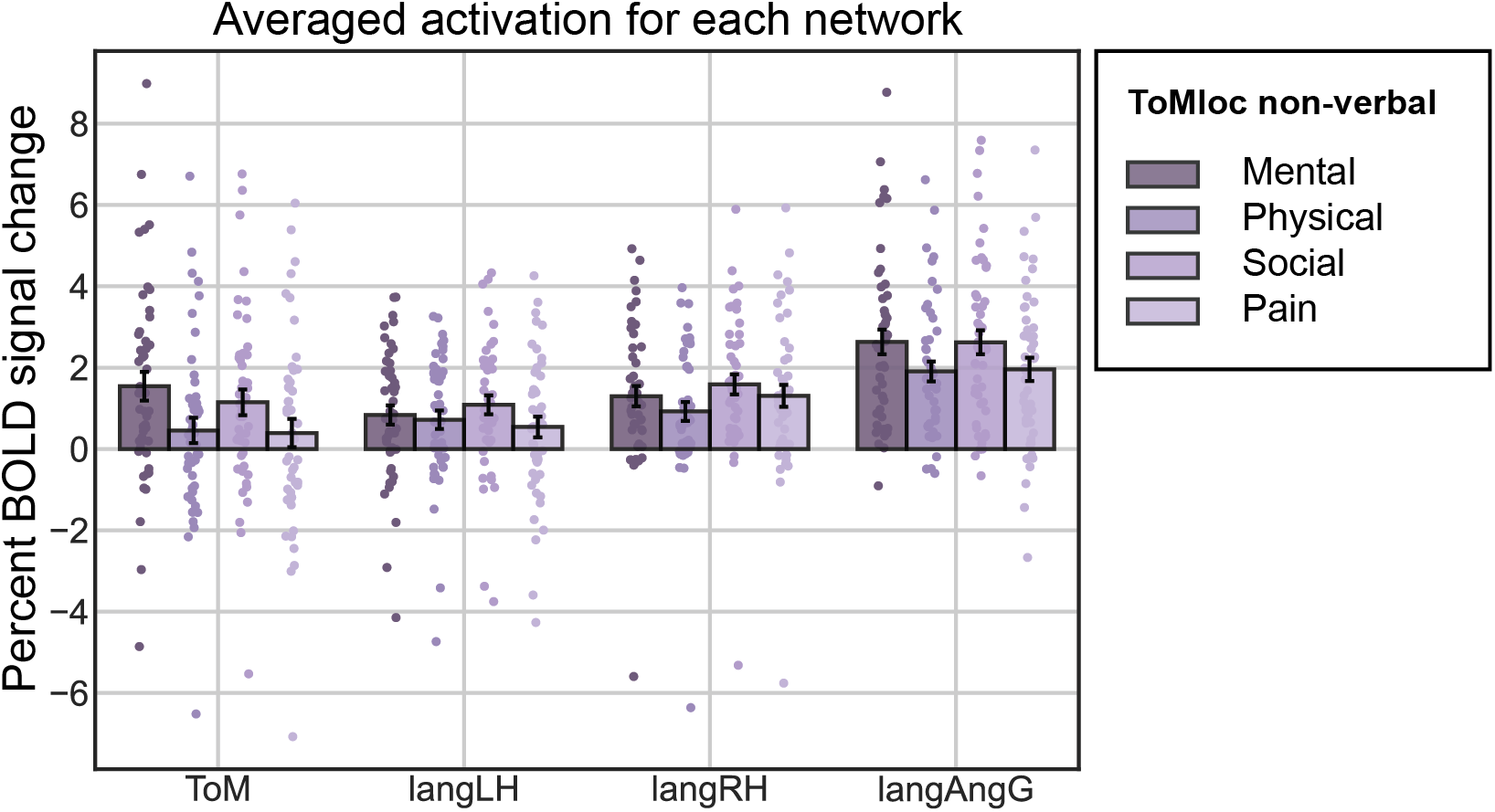
Responses by network to the four conditions of the non-verbal ToM localizer (Mental, Physical, Social, and Pain). Whereas the ToM network is most strongly engaged by the Mental condition, language regions in both hemispheres (langLH and langRH) are most strongly engaged by the Social condition, and thus, in contrast to the ToM network, neither shows a selective response to mentalizing (Mental > Social). Language selective regions of the bilateral angular gyri (langAngG) are systematically more engaged by social aspects of the task (Mental/Social > Physical/Pain) but likewise show no selectivity for mentalizing (Mental > Social).

Note that, based on much prior work (e.g., Saxe & Kanwisher, 2003; Saxe & Powell, 2006; Van Overwalle, 2009; for review see e.g., Saxe et al., 2004; Van Overwalle, 2009), we are assuming the existence of the ToM network (i.e. a brain network that selectively supports theory of mind reasoning and that is spatially and functionally distinct from the language periphery), and we are merely reporting its responses for reference. Nonetheless, a surprising finding in **Figure 8** is that the ToM network itself shows a relatively large response to the *social* condition, unlike prior studies that reported clearer selectivity for the *mental* condition (Jacoby et al., 2016), suggesting that areas identified by the verbal ToM localizer may show a more generalized social response, albeit weaker than the response to mental state content. Since our present focus is on the functional role of language areas in ToM, rather than on previously established ToM areas, we leave further investigation of these questions (i.e., the contributions of the ToM network to social functions beyond mentalizing, and the relationship between the ToM network and the language periphery) to future work.

## Discussion

Given the close functional relationship between language processing and thinking about others’ thoughts (theory of mind, ToM), both developmentally (e.g., Astington & Jenkins, 1999; Peterson & Siegal, 2000; Hale & Tager-Flusberg, 2003; Ruffman et al., 2003; Astington & Baird, 2005; Slade & Ruffman, 2005; Miller, 2006; de Villiers & de Villiers, 2014) and in adult language use (e.g., Grice, 1975; Sperber & Wilson, 1987; Winner et al., 1998; Champagne-Lavau & Joanette, 2009; Roberts, 2012), we asked whether the human language network, or some of its components, might additionally represent ToM information, as indicated by recent findings from Deen et al (2015). To investigate this question, we localized the language network in each participant in a large-scale fMRI study and evaluated the responses of these language areas to the established verbal ToM localizer task (based on the false belief > false photo contrast) and a more recently introduced nonverbal ToM task (based on the mental movie segments > physical movie segments contrast). Although the language network responds significantly to the verbal ToM contrast, it does not respond to the nonverbal ToM contrast, suggesting that the verbal ToM effect may be an artifact of linguistic differences between the conditions of the verbal ToM localizer. We confirmed this hypothesis by analyzing the verbal ToM materials with respect to linguistic features that are independently known to modulate activity in the language network. We showed that controlling for these features strongly attenuates the verbal ToM effect in language areas. It is thus likely that prior reports of language network activation in response to the verbal ToM contrast (Deen et al., 2015) were affected by these linguistic confounds.

In short, we do not find evidence that core language areas are engaged in ToM reasoning. Nonetheless, both the non-verbal mental > physical contrast and the linguistically residualized verbal false belief > false photo contrast are numerically positive in the language network as a whole as well as in each component fROI. We cannot rule out the possibility that these regions show a small increase in response to mentalizing that our current (relatively large) sample (n=149 for verbal ToM and n=48 for non-verbal ToM) lacks power to detect. However, we have shown that any such effects are much smaller than effects of language processing, and thus that the functional profile of these regions overwhelmingly favors language over ToM. Furthermore, the core language network does not show the characteristic selectivity for mentalizing; video segments depicting non-mentalizing social interactions induce a similar magnitude response to segments involving mentalizing. Thus, the language network shows neither a general response to ToM, nor selectivity for ToM relative to other kinds of social processing, and prior evidence to the contrary (e.g., Deen et al., 2015) may have been driven by linguistic confounds in the standard ToM localizer. Our results thus converge with recent findings from resting state functional correlation analyses that independently identify a ToM-selective *default network B* (DN-B; Braga & Buckner, 2017; DiNicola et al., 2020) and show that this network is spatially distinct from the language network in individual brains (Braga et al., 2020).

In addition to our critical question about the involvement of core language areas in ToM processing, we additionally a investigated possible role in ToM of areas in the “periphery” of the language network (Chai et al., 2016) that have been implicated by prior work in ToM, social processing, or social/affective aspects of language processing: the right hemisphere (RH) homotopes of core language areas, as well as language areas in the bilateral angular gyri.

The RH homotopes of the language regions respond to language contrasts, although generally less strongly (e.g., Fedorenko et al., 2010; Mahowald & Fedorenko, 2016; Quillen et al., 2021; Lipkin et al., in press; Martin et al., 2022). A number of claims have been made about the role of RH language regions in language processing and differential contributions of LH vs. RH language regions (e.g., Ross & Mesulam, 1979; Bryan, 1989; Bottini et al., 1994; Van Lancker, 1997; Mitchell & Crow, 2005; Lindell, 2006; Beeman & Chiarello, 2013). A common theme in this literature associates RH homotopes of language areas with social, pragmatic, nonliteral, and/or affective aspects of speech processing and/or language comprehension (e.g., van Lancker, 1997; Mitchell & Crow, 2005), including potentially a role in leveraging ToM for pragmatic inference (Kaplan et al., 1990). However, the empirical landscape is complex and ridden with controversy. Even the most common claim about the stronger role of the RH language areas, compared to the LH language areas, in non-literal comprehension has been questioned (e.g., Lee & Dapretto, 2006; Rapp et al., 2007, 2012; Paunov et al., 2019; Hauptman et al., 2022; see e.g., Calvo et al., 2019 for patient evidence). Based on analyses of data from the Human Connectome Project (Van Essen et al., 2013), Rajimehr et al. (2021) recently argued that the primary function of these areas may be social rather than linguistic.

The function of the language-responsive areas in the left and right angular gyri also remains debated. A number of proposals have been put forward about the angular gyri in general (e.g., Farrer et al., 2008; Bonner et al., 2013; Price et al., 2015; Davis & Yee, 2019; Humphreys et al., 2021) and their specific role in language processing (e.g., Thothathiri et al., 2012; Bemis & Pylkkänen, 2013; Matchin et al., 2019; Branzi et al., 2021) as well as ToM processing (e.g., Saxe & Kanwisher, 2003; Schurz et al., 2014, 2017). But, like other parts of the association cortex, the angular gyrus is a highly structurally and functionally heterogeneous area (e.g., Scholz et al., 2009; Uddin et al., 2010; Seghier, 2013), which makes proposals about the entire angular gyrus difficult to evaluate. Of most relevance to the current investigation, Deen et al. (2015) observed some overlap between linguistic and ToM contrasts at the individual-participant level in the angular gyrus.

Unlike the core LH language areas, our analyses of the language periphery revealed a robust mental > physical contrast in the non-verbal ToM localizer, indicating that the language periphery indeed responds to mental content across representational formats (verbal and visual). However, unlike the ToM network, language fROIs in the right hemisphere and in the bilateral angular gyri respond no less strongly to non-mentalizing social interactions, and the RH fROIs additionally register a strong response to observing others’ physical pain. These response characteristics are not consistent with a selective response to ToM in the language periphery. They could be consistent with a broadly social function as proposed by e.g., Rajimehr et al. (2021) for the RH language homotopes. However, Rajimehr et al.’s claim is based on a single paradigm evaluated in a single (albeit large) dataset, and alternative explanations in terms of, for example, general visual semantic processing (e.g., Zaidel, 1987; Joseph, 1988) cannot be ruled out. Under such accounts, the somewhat stronger responses to social conditions would be explained by greater overall attention to social content. Thus, more research is needed to understand the precise contribution of the RH language homotopes to semantic and specifically social cognition.

If the language network is not involved in making inferences about others’ thoughts, how then do these inferences enter into language processing in order to inform rapid incremental sentence comprehension (e.g., Shibata et al., 2010; Regel et al., 2011; Kaakinen et al., 2014)? We hypothesize that this occurs via rapid feedback from the ToM network, which can then be used to inform interpretation. Although further research is needed to investigate this hypothesis, prior work has shown that the language and ToM networks show reliable functional correlations with each other over time during naturalistic cognition, consistent with information sharing (Paunov et al., 2019).

In conclusion, fMRI evidence supports a spatial dissociation between core language processing areas on the one hand and areas involved in making inferences about others’ mental states (ToM). We find no evidence of mentalizing in the core LH language network using a non-verbal ToM task, and we further find no selectivity for mentalizing over other kinds of social cognition. Linguistic analyses indicate that prior reports of overlap between the language and ToM networks may have been driven by confounds with linguistic variables independently known to drive language network activity. These results do not support a role for the language network in making inferences about others’ mental states. The language “periphery”—consisting of the RH homotopic language areas and the language-responsive areas in the bilateral angular gyri—responds relatively more strongly than the core language network to conditions that encourage mentalizing. However, these stronger responses also extend to other kinds of social conditions and even non-social ones, consistent with these regions’ role in social and even general visual-semantic processing.

## Acknowledgments

We would like to thank Zach Mineroff, Melissa Kline, and Brianna Pritchett for helping with fMRI data collection; Rebecca Saxe and Ben Deen for comments on the earlier draft of the manuscript; and EvLab and TedLab members for helpful discussions. We would also like to acknowledge the Athinoula A. Martinos Imaging Center at the McGovern Institute for Brain Research at MIT, and its support team (Steve Shannon, and Atsushi Takahashi). This work was supported by the NIH award R01-DC016607. EF was additionally supported by the NIH award R01-DC016950, by a grant from the Simons Foundation to the Simons Center for the Social Brain at MIT, and by research funds from the McGovern Institute for Brain Research and the Department of Brain and Cognitive Sciences.

## References

Apperly, I. A., Samson, D., Carroll, N., Hussain, S., & Humphreys, G. (2006). Intact first-and second-order false belief reasoning in a patient with severely impaired grammar. Social Neuroscience, 1(3– 4), 334–348.

Apperly, I. A., Samson, D., Chiavarino, C., & Humphreys, G. W. (2004). Frontal and temporo-parietal lobe contributions to theory of mind: neuropsychological evidence from a false-belief task with reduced language and executive demands. Journal of Cognitive Neuroscience, 16(10), 1773–1784.

Astington, J. W., & Baird, J. A. (2005). Why language matters for theory of mind. Oxford University Press.

Astington, J. W., & Jenkins, J. M. (1999). A longitudinal study of the relation between language and theory-of-mind development. Developmental Psychology, 35(5), 1311.

Beeman, M. J., & Chiarello, C. (2013). Right hemisphere language comprehension: Perspectives from cognitive neuroscience. Psychology Press.

Bemis, D. K., & Pylkkänen, L. (2013). Basic linguistic composition recruits the left anterior temporal lobe and left angular gyrus during both listening and reading. Cerebral Cortex, 23(8), 1859–1873.

Benjamini, Y., & Yekutieli, D. (2001). The control of the false discovery rate in multiple testing under dependency. Annals of Statistics, 1165–1188.

Blank, I., Balewski, Z., Mahowald, K., & Fedorenko, E. (2016). Syntactic processing is distributed across the language system. Neuroimage, 127, 307–323.

Blank, I., Kanwisher, N., & Fedorenko, E. (2014). A functional dissociation between language and multiple-demand systems revealed in patterns of BOLD signal fluctuations. Journal of Neurophysiology, 112(5), 1105–1118.

Bonner, M. F., Peelle, J. E., Cook, P. A., & Grossman, M. (2013). Heteromodal conceptual processing in the angular gyrus. Neuroimage, 71, 175–186.

Bottini, G., Corcoran, R., Sterzi, R., Paulesu, E., Schenone, P., Scarpa, P., Frackowiak, R. S. J., & Frith, D. (1994). The role of the right hemisphere in the interpretation of figurative aspects of language A positron emission tomography activation study. Brain, 117(6), 1241–1253.

Braga, R. M., & Buckner, R. L. (2017). Parallel interdigitated distributed networks within the individual estimated by intrinsic functional connectivity. Neuron, 95(2), 457–471.

Braga, R. M., DiNicola, L. M., Becker, H. C., & Buckner, R. L. (2020). Situating the left-lateralized language network in the broader organization of multiple specialized large-scale distributed networks. Journal of Neurophysiology, 124(5), 1415–1448.

Branzi, F. M., Pobric, G., Jung, J., & Lambon Ralph, M. A. (2021). The left angular gyrus is causally involved in context-dependent integration and associative encoding during narrative reading. Journal of Cognitive Neuroscience, 33(6), 1082–1095.

Brothers, L., & Ring, B. (1992). A neuroethological framework for the representation of minds. Journal of Cognitive Neuroscience, 4(2), 107–118.

Bruneau, E. G., Dufour, N., & Saxe, R. (2012). Social cognition in members of conflict groups: behavioural and neural responses in Arabs, Israelis and South Americans to each other’s misfortunes. Philosophical Transactions of the Royal Society B: Biological Sciences, 367(1589), 717–730.

Bruneau, E. G., Pluta, A., & Saxe, R. (2012). Distinct roles of the ‘shared pain’and ‘theory of mind’networks in processing others’ emotional suffering. Neuropsychologia, 50(2), 219–231.

Bryan, K. L. (1989). Language prosody and the right hemisphere. Aphasiology, 3(4), 285–299.

Calvo, N., Abrevaya, S., Martínez Cuitiño, M., Steeb, B., Zamora, D., Sedeño, L., Ibáñez, A., & García, A. M. (2019). Rethinking the Neural Basis of Prosody and Non-literal Language: Spared Pragmatics and Cognitive Compensation in a Bilingual With Extensive Right-Hemisphere Damage. Frontiers in Psychology, 10. https://doi.org/10.3389/fpsyg.2019.00570

Castelli, F., Frith, C., Happé, F., & Frith, U. (2002). Autism, Asperger syndrome and brain mechanisms for the attribution of mental states to animated shapes. Brain, 125(8), 1839–1849.

Castelli, F., Happé, F., Frith, U., & Frith, C. (2000). Movement and mind: A functional imaging study of perception and interpretation of complex intentional movement patterns. NeuroImage, 12(3), 314– 325.

Chai, L. R., Mattar, M. G., Blank, I. A., Fedorenko, E., & Bassett, D. S. (2016). Functional network dynamics of the language system. Cerebral Cortex, 26(11), 4148–4159.

Champagne-Lavau, M., & Joanette, Y. (2009). Pragmatics, theory of mind and executive functions after a right-hemisphere lesion: Different patterns of deficits. Journal of Neurolinguistics, 22(5), 413–426.

Davis, C. P., & Yee, E. (2019). Features, labels, space, and time: Factors supporting taxonomic relationships in the anterior temporal lobe and thematic relationships in the angular gyrus. Language, Cognition and Neuroscience, 34(10), 1347–1357.

de Villiers, J. G., & de Villiers, P. A. (2014). The role of language in theory of mind development. Topics in Language Disorders, 34(4), 313–328.

Deen, B., Koldewyn, K., Kanwisher, N., & Saxe, R. (2015). Functional organization of social perception and cognition in the superior temporal sulcus. Cerebral Cortex, 25(11), 4596–4609.

Dennis, M., Simic, N., Bigler, E. D., Abildskov, T., Agostino, A., Taylor, H. G., Rubin, K., Vannatta, K., Gerhardt, C. A., Stancin, T., & others. (2013). Cognitive, affective, and conative theory of mind (ToM) in children with traumatic brain injury. Developmental Cognitive Neuroscience, 5, 25–39.

Diehl, J. J., Bennetto, L., & Young, E. C. (2006). Story recall and narrative coherence of high-functioning children with autism spectrum disorders. Journal of Abnormal Child Psychology, 34(1), 83–98.

DiNicola, L. M., Braga, R. M., & Buckner, R. L. (2020). Parallel distributed networks dissociate episodic and social functions within the individual. Journal of Neurophysiology, 123(3), 1144–1179.

Dodell-Feder, D., Koster-Hale, J., Bedny, M., & Saxe, R. (2011). fMRI item analysis in a theory of mind task. Neuroimage, 55(2), 705–712.

Domínguez, D. J. F., Nott, Z., Horne, K., Prangley, T., Adams, A. G., Henry, J. D., & Molenberghs, P. (2019). Structural and functional brain correlates of theory of mind impairment post-stroke. Cortex, 121, 427–442.

Dronkers, N. F., Ludy, C. A., & Redfern, B. B. (1998). Pragmatics in the absence of verbal language: Descriptions of a severe aphasic and a language-deprived adult. Journal of Neurolinguistics, 11(1–2), 179–190.

Dufour, N., Redcay, E., Young, L., Mavros, P. L., Moran, J. M., Triantafyllou, C., Gabrieli, J. D. E., & Saxe, R. (2013). Similar brain activation during false belief tasks in a large sample of adults with and without autism. PloS One, 8(9), e75468.

Farrer, C., Frey, S. H., Van Horn, J. D., Tunik, E., Turk, D., Inati, S., & Grafton, S. T. (2008). The angular gyrus computes action awareness representations. Cerebral Cortex, 18(2), 254–261.

Fedorenko, E., Behr, M. K., & Kanwisher, N. (2011). Functional specificity for high-level linguistic processing in the human brain. Proceedings of the National Academy of Sciences.

Fedorenko, E., Blank, I., Siegelman, M., & Mineroff, Z. (2020). Lack of selectivity for syntax relative to word meanings throughout the language network. Cognition, 203, 104348.

Fedorenko, E., Hsieh, P.-J., Nieto-Castañón, A., Whitfield-Gabrieli, S., & Kanwisher, N. (2010). New method for fMRI investigations of language: defining ROIs functionally in individual subjects. Journal of Neurophysiology, 104(2), 1177–1194.

Fedorenko, E., Scott, T. L., Brunner, P., Coon, W. G., Pritchett, B., Schalk, G., & Kanwisher, N. (2016). Neural correlate of the construction of sentence meaning. Proceedings of the National Academy of Sciences, 113(41), E6256--E6262.

Fedorenko, E., & Thompson-Schill, S. L. (2014). Reworking the language network. Trends in Cognitive Sciences, 18(3), 120–126.

Fletcher, P. C., Happe, F., Frith, U., Baker, S. C., Dolan, R. J., Frackowiak, R. S. J., & Frith, C. D. (1995). Other minds in the brain: a functional imaging study of “theory of mind” in story comprehension. Cognition, 57(2), 109–128.

Gallagher, H. L., Happé, F., Brunswick, N., Fletcher, P. C., Frith, U., & Frith, C. D. (2000). Reading the mind in cartoons and stories: an fMRI study of ‘theory of mind’in verbal and nonverbal tasks. Neuropsychologia, 38(1), 11–21.

Gibson, E. (2000). The Dependency Locality Theory: A distance-based theory of linguistic complexity. In A. Marantz, Y. Miyashita, & W. O’Neil (Eds.), Image, language, brain (pp. 95–106). MIT Press.

Graff, D., Kong, J., Chen, K., & Maeda, K. (2007). English Gigaword Third Edition LDC2007T07. Linguistic Data Consortium. https://catalog.ldc.upenn.edu/LDC2007T07

Grice, H. P. (1975). Logic and conversation. In P. Cole & J. L. Morgan (Eds.), Syntax and Semantics, Vol. 3: Speech Acts (pp. 41–58). Academic Press.

Hale, C. M., & Tager-Flusberg, H. (2003). The influence of language on theory of mind: A training study. Developmental Science, 6(3), 346–359.

Hauptman, M., Blank, I., & Fedorenko, E. (2022). Non-literal language processing is jointly supported by the language and Theory of Mind networks: Evidence from a novel meta-analytic fMRI approach. BioRxiv.

Heafield, K., Pouzyrevsky, I., Clark, J. H., & Koehn, P. (2013). Scalable modified Kneser-Ney language model estimation. Proceedings of the 51st Annual Meeting of the Association for Computational Linguistics, 690–696.

Hein, G., & Singer, T. (2008). I feel how you feel but not always: the empathic brain and its modulation. Current Opinion in Neurobiology, 18(2), 153–158.

Humphreys, G. F., Ralph, M. A. L., & Simons, J. S. (2021). A unifying account of angular gyrus contributions to episodic and semantic cognition. Trends in Neurosciences, 44(6), 452–463.

Jacoby, N., Bruneau, E., Koster-Hale, J., & Saxe, R. (2016). Localizing Pain Matrix and Theory of Mind networks with both verbal and non-verbal stimuli. Neuroimage, 126, 39–48.

Joseph, R. (1988). The right cerebral hemisphere: Emotion, music, visual-spatial skills, body-image, dreams, and awareness. Journal of Clinical Psychology, 44(5), 630–673.

Just, M. A., & Carpenter, P. A. (1980). A theory of reading: From eye fixations to comprehension. Psychological Review, 87(4), 329–354.

Kaakinen, J. K., Olkoniemi, H., Kinnari, T., & Hyönä, J. (2014). Processing of written irony: An eye movement study. Discourse Processes, 51(4), 287–311.

Kamps, F. S., Richardson, H., Murty, N. A. R., Kanwisher, N., & Saxe, R. (2022). Using child-friendly movie stimuli to study the development of face, place, and object regions from age 3 to 12 years. Human Brain Mapping. https://doi.org/https://doi.org/10.1002/hbm.25815

Kaplan, J. A., Brownell, H. H., Jacobs, J. R., & Gardner, H. (1990). The effects of right hemisphere damage on the pragmatic interpretation of conversational remarks. Brain and Language, 38(2), 315– 333.

Koster-Hale, J., & Saxe, R. (2013). Theory of mind: A neural prediction problem. Neuron, 79(5), 836– 848.

Kriegeskorte, N., Simmons, W. K., Bellgowan, P. S. F., & Baker, C. I. (2009). Circular analysis in systems neuroscience: The dangers of double dipping. Nature Neuroscience, 12(5), 535–540.

Lee, S. S., & Dapretto, M. (2006). Metaphorical vs. literal word meanings: fMRI evidence against a selective role of the right hemisphere. NeuroImage, 29(2), 536–544.

Lindell, A. K. (2006). In your right mind: Right hemisphere contributions to language processing and production. Neuropsychology Review, 16(3), 131–148.

Lipkin, B., Tuckute, G., Affourtit, J., Small, H., Mineroff, Z., Kean, H., Jouravlev, O., Rakocevic, L., Pritchett, B., Siegelman, M., & others. (2022). LanA (Language Atlas): A probabilistic atlas for the language network based on fMRI data from > 800 individuals. BioRxiv.

Lombardo, M. V, Chakrabarti, B., Bullmore, E. T., Baron-Cohen, S., Consortium, M. R. C. A., & others. (2011). Specialization of right temporo-parietal junction for mentalizing and its relation to social impairments in autism. Neuroimage, 56(3), 1832–1838.

Lopopolo, A., Frank, S. L., den Bosch, A., & Willems, R. M. (2017). Using stochastic language models (SLM) to map lexical, syntactic, and phonological information processing in the brain. PloS One, 12(5), e0177794.

Malik-Moraleda, S., Ayyash, D., Gallée, J., Affourtit, J., Hoffman, M., Mineroff, Z., Jouravlev, O., & Fedorenko, E. (2022). The universal language network: A cross-linguistic investigation spanning 45 languages and 11 language families. BioRxiv.

Mar, R. A. (2011). The neural bases of social cognition and story comprehension. Annual Review of Psychology, 62, 103–134.

Marcus, M. P., Santorini, B., & Marcinkiewicz, M. A. (1993). Building a large annotated corpus of English: the Penn Treebank. Computational Linguistics, 19(2), 313–330.

Martín-Rodríguez, J. F., & León-Carrión, J. (2010). Theory of mind deficits in patients with acquired brain injury: A quantitative review. Neuropsychologia, 48(5), 1181–1191.

Martin, K. C., Seydell-Greenwald, A., Berl, M. M., Gaillard, W. D., Turkeltaub, P. E., & Newport, E. L. (2022). A weak shadow of early life language processing persists in the right hemisphere of the mature brain. Neurobiology of Language, 1–49.

Mason, R. A., & Just, M. A. (2011). Differentiable cortical networks for inferences concerning people’s intentions versus physical causality. Human Brain Mapping, 32(2), 313–329.

Matchin, W., Liao, C.-H., Gaston, P., & Lau, E. (2019). Same words, different structures: An fMRI investigation of argument relations and the angular gyrus. Neuropsychologia, 125, 116–128.

Miller, C. A. (2006). Developmental relationships between language and theory of mind.

Mitchell, R. L. C., & Crow, T. J. (2005). Right hemisphere language functions and schizophrenia: the forgotten hemisphere? Brain, 128(5), 963–978.

Nelson, M. J., El Karoui, I., Giber, K., Yang, X., Cohen, L., Koopman, H., Cash, S. S., Naccache, L., Hale, J. T., Pallier, C., & others. (2017). Neurophysiological dynamics of phrase-structure building during sentence processing. Proceedings of the National Academy of Sciences, 114(18), E3669--E3678.

Nguyen, L., van Schijndel, M., & Schuler, W. (2012). Accurate Unbounded Dependency Recovery using Generalized Categorial Grammars. Proceedings of COLING 2012.

Nieto-Castañón, A., & Fedorenko, E. (2012). Subject-specific functional localizers increase sensitivity and functional resolution of multi-subject analyses. Neuroimage, 63(3), 1646–1669.

Oldfield, R. C. (1971). The assessment and analysis of handedness: the Edinburgh inventory. Neuropsychologia, 9(1), 97–113.

Pallier, C., Devauchelle, A.-D., & Dehaene, S. (2011). Cortical representation of the constituent structure of sentences. Proceedings of the National Academy of Sciences, 108(6), 2522–2527.

Paunov, A., Blank, I. A., Jouravlev, O., Mineroff, Z., Gallée, J., & Fedorenko, E. (2022). Differential tracking of linguistic vs. mental state content in naturalistic stimuli by language and Theory of Mind (ToM) brain networks. BioRxiv.

Paunov, A., Blank, I., & Fedorenko, E. (2019). Functionally distinct language and Theory of Mind networks are synchronized at rest and during language comprehension. Journal of Neurophysiology, 121(4), 1244–1265. https://doi.org/10.1152/jn.00619.2018

Peterson, C. C., & Siegal, M. (2000). Insights into theory of mind from deafness and autism. Mind & Language, 15(1), 123–145.

Price, A. R., Bonner, M. F., Peelle, J. E., & Grossman, M. (2015). Converging evidence for the neuroanatomic basis of combinatorial semantics in the angular gyrus. Journal of Neuroscience, 35(7), 3276–3284.

Pritchett, B. L., Hoeflin, C., Koldewyn, K., Dechter, E., & Fedorenko, E. (2018). High-level language processing regions are not engaged in action observation or imitation. Journal of Neurophysiology, 120(5), 2555–2570.

Quillen, I. A., Yen, M., & Wilson, S. M. (2021). Distinct Neural Correlates of Linguistic and Non-Linguistic Demand. Neurobiology of Language, 2(2), 202–225.

Rajimehr, R., Firoozi, A., Rafipoor, H., Abbasi, N., & Duncan, J. (2021). Complementary hemispheric lateralization of language and social processing in the human brain.

Rapp, A. M., Leube, D. T., Erb, M., Grodd, W., & Kircher, T. T. J. (2007). Laterality in metaphor processing: Lack of evidence from functional magnetic resonance imaging for the right hemisphere theory. Brain and Language, 100(2), 142–149.

Rapp, A. M., Mutschler, D. E., & Erb, M. (2012). Where in the brain is nonliteral language? A coordinate-based meta-analysis of functional magnetic resonance imaging studies. Neuroimage, 63(1), 600–610.

Rayner, K., Kambe, G., & Duffy, S. A. (2000). The effect of clause wrap-up on eye movements during reading. The Quarterly Journal of Experimental Psychology: Section A, 53(4), 1061–1080.

Regel, S., Gunter, T. C., & Friederici, A. D. (2011). Isn’t it ironic? An electrophysiological exploration of figurative language processing. Journal of Cognitive Neuroscience, 23(2), 277–293.

Richardson, H., Koster-Hale, J., Caselli, N., Magid, R., Benedict, R., Olson, H., Pyers, J., & Saxe, R. (2020). Reduced neural selectivity for mental states in deaf children with delayed exposure to sign language. Nature Communications, 11(1), 1–13.

Richardson, H., Lisandrelli, G., Riobueno-Naylor, A., & Saxe, R. (2018). Development of the social brain from age three to twelve years. Nature Communications, 9(1), 1–12.

Roberts, C. (2012). Information structure: Towards an integrated formal theory of pragmatics. Semantics and Pragmatics, 5, 1–6.

Ross, E. D., & Mesulam, M.-M. (1979). Dominant language functions of the right hemisphere?: Prosody and emotional gesturing. Archives of Neurology, 36(3), 144–148.

Ruby, P., & Decety, J. (2003). What you believe versus what you think they believe: A neuroimaging study of conceptual perspective-taking. European Journal of Neuroscience, 17(11), 2475–2480.

Ruffman, T., Slade, L., Rowlandson, K., Rumsey, C., & Garnham, A. (2003). How language relates to belief, desire, and emotion understanding. Cognitive Development, 18(2), 139–158.

Saxe, R. (2006). Uniquely human social cognition. Current Opinion in Neurobiology, 16(2), 235–239.

Saxe, R. (2010). The right temporo-parietal junction: a specific brain region for thinking about thoughts. Handbook of Theory of Mind, 1–35.

Saxe, R., Brett, M., & Kanwisher, N. (2006). Divide and conquer: a defense of functional localizers. Neuroimage, 30(4), 1088–1096.

Saxe, R., Carey, S., & Kanwisher, N. (2004). Understanding other minds. Annu. Rev. Psychol, 55, 87– 124.

Saxe, R., & Kanwisher, N. (2003). People thinking about thinking people: The role of the temporo-parietal junction in “theory of mind.” Neuroimage, 19(4), 1835–1842.

Saxe, R., & Powell, L. J. (2006). It’s the thought that counts: specific brain regions for one component of theory of mind. Psychological Science, 17(8), 692–699.

Saxe, R., & Wexler, A. (2005). Making sense of another mind: the role of the right temporo-parietal junction. Neuropsychologia, 43(10), 1391–1399.

Scholz, J., Triantafyllou, C., Whitfield-Gabrieli, S., Brown, E. N., & Saxe, R. (2009). Distinct regions of right temporo-parietal junction are selective for theory of mind and exogenous attention. PloS One, 4(3), e4869.

Schurz, M., Radua, J., Aichhorn, M., Richlan, F., & Perner, J. (2014). Fractionating theory of mind: a meta-analysis of functional brain imaging studies. Neuroscience & Biobehavioral Reviews, 42, 9– 34.

Schurz, M., Tholen, M. G., Perner, J., Mars, R. B., & Sallet, J. (2017). Specifying the brain anatomy underlying temporo-parietal junction activations for theory of mind: A review using probabilistic atlases from different imaging modalities. Human Brain Mapping, 38(9), 4788–4805.

Schuster, S., Hawelka, S., Hutzler, F., Kronbichler, M., & Richlan, F. (2016). Words in context: The effects of length, frequency, and predictability on brain responses during natural reading. Cerebral Cortex, 26(10), 3889–3904.

Scott, T. L., Gallée, J., & Fedorenko, E. (2017). A new fun and robust version of an fMRI localizer for the frontotemporal language system. Cognitive Neuroscience, 8(3), 167–176.

Seghier, M. L. (2013). The angular gyrus: multiple functions and multiple subdivisions. The Neuroscientist, 19(1), 43–61.

Shain, C. (2019). A large-scale study of the effects of word frequency and predictability in naturalistic reading. Proceedings of the 2019 Conference of the North American Chapter of the Association for Computational Linguistics: Human Language Technologies, Volume 1 (Long and Short Papers), 4086–4094.

Shain, C., Blank, I. A., Fedorenko, E., Gibson, E., & Schuler, W. (2021). Robust effects of working memory demand during naturalistic language comprehension in language-selective cortex. BioRxiv. https://doi.org/10.1101/2021.09.18.460917

Shain, C., Blank, I., van Schijndel, M., Schuler, W., & Fedorenko, E. (2020). fMRI reveals language-specific predictive coding during naturalistic sentence comprehension. Neuropsychologia, 138, 107307.

Shamay-Tsoory, S. G., Harari, H., Aharon-Peretz, J., & Levkovitz, Y. (2010). The role of the orbitofrontal cortex in affective theory of mind deficits in criminal offenders with psychopathic tendencies. Cortex, 46(5), 668–677.

Shibata, M., Toyomura, A., Itoh, H., & Abe, J. (2010). Neural substrates of irony comprehension: A functional MRI study. Brain Research, 1308, 114–123.

Singer, T., & Lamm, C. (2009). The social neuroscience of empathy. Annals of the New York Academy of Sciences, 1156(1), 81–96.

Slade, L., & Ruffman, T. (2005). How language does (and does not) relate to theory of mind: A longitudinal study of syntax, semantics, working memory and false belief. British Journal of Developmental Psychology, 23(1), 117–141.

Small, H., Lipkin, B., Affourtit, J., Pongos, A., & Fedorenko, E. (2021). Differential selectivity of the left and right hemisphere language regions for non-linguistic processing. Proceedings of the Thirteenth Annual Meeting of the Society for the Neurobiology of Language, 264.

Sommer, M., Döhnel, K., Sodian, B., Meinhardt, J., Thoermer, C., & Hajak, G. (2007). Neural correlates of true and false belief reasoning. Neuroimage, 35(3), 1378–1384.

Sperber, D., & Wilson, D. (1987). Précis of relevance: Communication and cognition. Behavioral and Brain Sciences, 10(4), 697–710.

Sprong, M., Schothorst, P., Vos, E., Hox, J., & Van Engeland, H. (2007). Theory of mind in schizophrenia: meta-analysis. The British Journal of Psychiatry, 191(1), 5–13.

Tager-Flusberg, H., Paul, R., & Lord, C. (2005). Language and Communication in Autism. Handbook of Autism and Pervasive Developmental Disorders, 1, 335–364.

Thothathiri, M., Kimberg, D. Y., & Schwartz, M. F. (2012). The neural basis of reversible sentence comprehension: evidence from voxel-based lesion symptom mapping in aphasia. Journal of Cognitive Neuroscience, 24(1), 212–222.

Uddin, L. Q., Supekar, K., Amin, H., Rykhlevskaia, E., Nguyen, D. A., Greicius, M. D., & Menon, V. (2010). Dissociable connectivity within human angular gyrus and intraparietal sulcus: evidence from functional and structural connectivity. Cerebral Cortex, 20(11), 2636–2646.

Van Essen, D. C., Smith, S. M., Barch, D. M., Behrens, T. E. J., Yacoub, E., Ugurbil, K., Wu-Minn H C P Consortium, & others. (2013). The WU-Minn human connectome project: an overview. Neuroimage, 80, 62–79.

Van Lancker, D. (1997). Rags to riches: our increasing appreciation of cognitive and communicative abilities of the human right cerebral hemisphere. Brain and Language, 57(1), 1–11.

Van Overwalle, F. (2009). Social cognition and the brain: a meta-analysis. Human Brain Mapping, 30(3), 829–858.

van Schijndel, M., Exley, A., & Schuler, W. (2013). A model of language processing as hierarchic sequential prediction. Topics in Cognitive Science, 5(3), 522–540. https://doi.org/10.1111/tops.12034

Varley, R., Siegal, M., & Want, S. C. (2001). Severe impairment in grammar does not preclude theory of mind. Neurocase, 7(6), 489–493.

Vogeley, K., Bussfeld, P., Newen, A., Herrmann, S., Happé, F., Falkai, P., Maier, W., Shah, N. J., Fink, G. R., & Zilles, K. (2001). Mind reading: neural mechanisms of theory of mind and self-perspective. Neuroimage, 14(1), 170–181.

Wellman, H. M., Cross, D., & Watson, J. (2001). Meta-analysis of theory-of-mind development: The truth about false belief. Child Development, 72(3), 655–684.

Willems, R. M., Benn, Y., Hagoort, P., Toni, I., & Varley, R. (2011). Communicating without a functioning language system: implications for the role of language in mentalizing. Neuropsychologia, 49(11), 3130–3135.

Willems, R. M., der Haegen, L., Fisher, S. E., & Francks, C. (2014). On the other hand: Including left-handers in cognitive neuroscience and neurogenetics. Nature Reviews Neuroscience, 15(3), 193.

Wimmer, H., & Perner, J. (1983). Beliefs about beliefs: Representation and constraining function of wrong beliefs in young children’s understanding of deception. Cognition, 13(1), 103–128.

Winner, E., Brownell, H., Happé, F., Blum, A., & Pincus, D. (1998). Distinguishing lies from jokes: Theory of mind deficits and discourse interpretation in right hemisphere brain-damaged patients. Brain and Language, 62(1), 89–106.

Young, L., Dodell-Feder, D., & Saxe, R. (2010). What gets the attention of the temporo-parietal junction? An fMRI investigation of attention and theory of mind. Neuropsychologia, 48(9), 2658–2664.

Zaidel, D. W. (1987). Hemispheric asymmetry in long-term semantic relationships. Cognitive Neuropsychology, 4(3), 321–332.

